# Estimating heritability and its enrichment in tissue-specific gene sets in admixed populations

**DOI:** 10.1101/503144

**Authors:** Yang Luo, Xinyi Li, Xin Wang, Steven Gazal, Josep Maria Mercader, 23andMe Research Team, SIGMA Type 2 Diabetes Consortium, Benjamin M. Neale, Jose C. Florez, Adam Auton, Alkes L. Price, Hilary K. Finucane, Soumya Raychaudhuri

**Author notes:** These authors contributed equally to this work (Yang Luo, Xinyi Li). These authors contributed equally to this work (Hilary K. Finucane, Soumya Raychaudhuri). Correspondence should be addressed to H.K.F. or S.R.

## Abstract

The increasing size and diversity of genome-wide association studies provide an exciting opportunity to study how the genetics of complex traits vary among diverse populations. Here, we introduce covariate-adjusted LD score regression (cov-LDSC), a method to accurately estimate genetic heritability 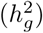 and its enrichment in both homogenous and admixed populations with summary statistics and in-sample LD estimates. In-sample LD can be estimated from a subset of the GWAS samples, allowing our method to be applied efficiently to very large cohorts. In simulations, we show that unadjusted LDSC underestimates 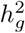 by 10% − 60% in admixed populations; in contrast, cov-LDSC is robust to all simulation parameters. We apply cov-LDSC to genotyping data from approximately 170,000 Latino, 47,000 African American and 135,000 European individuals. We estimate 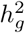 and detect heritability enrichment in three quantitative and five dichotomous phenotypes respectively, making this, to our knowledge, the most comprehensive heritability-based analysis of admixed individuals. Our results show that most traits have high concordance of 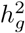 and consistent tissue-specific heritability enrichment among different populations. However, for age at menarche, we observe population-specific heritability estimates of 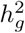. We observe consistent patterns of tissue-specific heritability enrichment across populations; for example, in the limbic system for BMI, the per-standardized-annotation effect size *τ*\* is 0.16 ± 0.04, 0.28 ± 0.11 and 0.18 ± 0.03 in Latino, African American and European populations respectively. Our results demonstrate that our approach is a powerful way to analyze genetic data for complex traits from underrepresented populations.

**Author summary:** Admixed populations such as African Americans and Hispanic Americans bear a disproportionately high burden of disease but remain underrepresented in current genetic studies. It is important to extend current methodological advancements for understanding the genetic basis of complex traits in homogeneous populations to individuals with admixed genetic backgrounds. Here, we develop a computationally efficient method to answer two specific questions. First, does genetic variation contribute to the same amount of phenotypic variation (heritability) across diverse populations? Second, are the genetic mechanisms shared among different populations? To answer these questions, we use our novel method to conduct the first comprehensive heritability-based analysis of a large number of admixed individuals. We show that there is a high degree of concordance in total heritability and tissue-specific enrichment between different ancestral groups. However, traits such as age at menarche show a noticeable differences among populations. Our work provides a powerful way to analyze genetic data in admixed populations and may contribute to the applicability of genomic medicine to admixed population groups.

## Introduction

It is important for human geneticists to study how genetic variants that influence phenotypic variability act across different populations worldwide [1, 2]. With increasingly large and diverse genetic studies, it is now becoming feasible to assess how the genetic mechanisms of complex traits act across populations. However, to date, most genome-wide association studies (GWAS) have been focused on relatively homogenous continental populations, and in particular those of European descent [3]. Non-European populations, particularly those with mixed ancestral backgrounds such as African Americans and Latinos, have been underrepresented in genetic studies. Many statistical methods to analyze genetic data assume homogeneous populations. In order to ensure that the benefits of GWAS are shared beyond individuals of homogeneous continental ancestry, statistical methods for admixed populations are needed [4].

Among methods to analyze polygenic complex traits in homogeneous populations, summary statistics-based methods such as linkage disequilibrium score regression (LDSC) [5, 6] and its extensions [7–9] have become particularly popular due to their computational efficiency, relative ease of application, and their applicability without raw genotyping data [10]. These methods can be used to estimate SNP-heritability, the proportion of phenotypic variance explained by genotyped variants [5, 11–13], distinguish polygenicity from confounding [5], establish relationships between complex phenotypes [7], and model genome-wide polygenic signals to identify key cell types and regulatory mechanisms of human diseases [6, 9, 14].

Summary statistics-based methods for polygenic analysis frequently rely on linkage disequilibrium (LD) calculations. For LD score regression, the LD information needed is the LD score for each SNP, defined to be the sum of its pairwise correlations (*r*^2^) with all other SNPs. For homogeneous populations there is usually a reference panel of individuals with matching ancestry that can be used to approximate the in-sample LD. For studies with heterogeneous or admixed ancestry, however, even when reference panels are available, they may not be representative of the precise populations used in the genetic study. For example Latino populations in different regions worldwide may share the same ancestral continental populations, but with dramatic differences in admixture proportions and timing of the admixture event [15]. A generic reference panel cannot easily capture these differences and hence cannot produce accurate LD scores that can be widely used for all Latino populations. Moreover, the structure of LD in heterogenous and admixed populations is complex and includes longer-range correlations that are absent or negligible in homogeneous populations. Thus, while LD scores computed from a matching reference panel reflect the appropriate matching LD for summary statistics computed in a homogeneous population, it has not been clear what the appropriate matching LD is for summary statistics computed in a heterogenous or admixed population, and so LDSC has only been recommended to be applied in homogeneous populations.

Here, we evaluate the heritability estimates using LDSC in admixed population and observe systematic underestimation. We then introduce covariate-adjusted LD score regression (cov-LDSC) to estimate heritability and partitioned heritability in admixed populations. We apply our approach to 8, 124 Latinos from a type 2 diabetes study (the Slim Initiative in Genomic Medicine for the Americas, SIGMA) [16] as well as 161, 894 Latino, 46, 844 African American, and 134, 999 European research participants from a personal genetics company (23andMe). We analyze three quantitative phenotypes (body mass index (BMI), height, and age at menarche), and five dichotomous phenotypes (type 2 diabetes (available in the SIGMA cohort only), left handedness, morning person, motion sickness, and nearsightedness).

One powerful component of LDSC is that it can be used to test whether a particular genome annotation -- for example, sets of genes that are specifically expressed within a candidate tissue or cell type -- capture more heritability than expected by chance [9, 11]. We demonstrate that cov-LDSC can be applied in the same way to identify trait-relevant tissue and cell types in admixed and homogenous populations with well-calibrated type I error. We examine height, BMI and morning person since these traits had sufficient statistical power [6] for cell-type enrichment analyses in the 23andMe cohort. We observe a high level of consistency among enriched tissue types, highlighting that the underlying biological processes are shared among studied populations. This heritability enrichment analysis of hundreds of genome annotations in cohorts of over 100,000 individuals would have been challenging with existing genotype-based methods [17–19].

## Results

### Overview of methods

In this work, we extended the LDSC-based methods to heterogeneous and admixed populations by introducing covariate-adjusted LDSC (cov-LDSC). We first showed through derivations that the appropriate matching LD for summary statistics computed in a heterogeneous or admixed population is in-sample LD computed on genotypes that have been adjusted for the same covariates (e.g. principal components) included in the summary statistics (S1 Appendix). In cov-LDSC, we compute these covariate-adjusted LD scores and then use LDSC to estimate heritability and its enrichment (**Methods**). We showed that, unlike LDSC, cov-LDSC produces accurate estimates of heritability with summary statistics from admixed populations (**Methods, Fig 1**). Furthermore, heritability can be partitioned to identify key gene sets that have disproportionately high heritability. While access to the genotype data of the GWAS samples is required to compute the covariate-adjusted LD scores, LD can be estimated on a random subset of the individuals, preserving the computational efficiency of LDSC and allowing for its application to very large studies. Individual cohorts can also release the in-sample covariate-adjusted LD scores as well as the summary statistics to avoid privacy concerns associated with genotype-level information to facilitate future studies.

**Fig 1.**
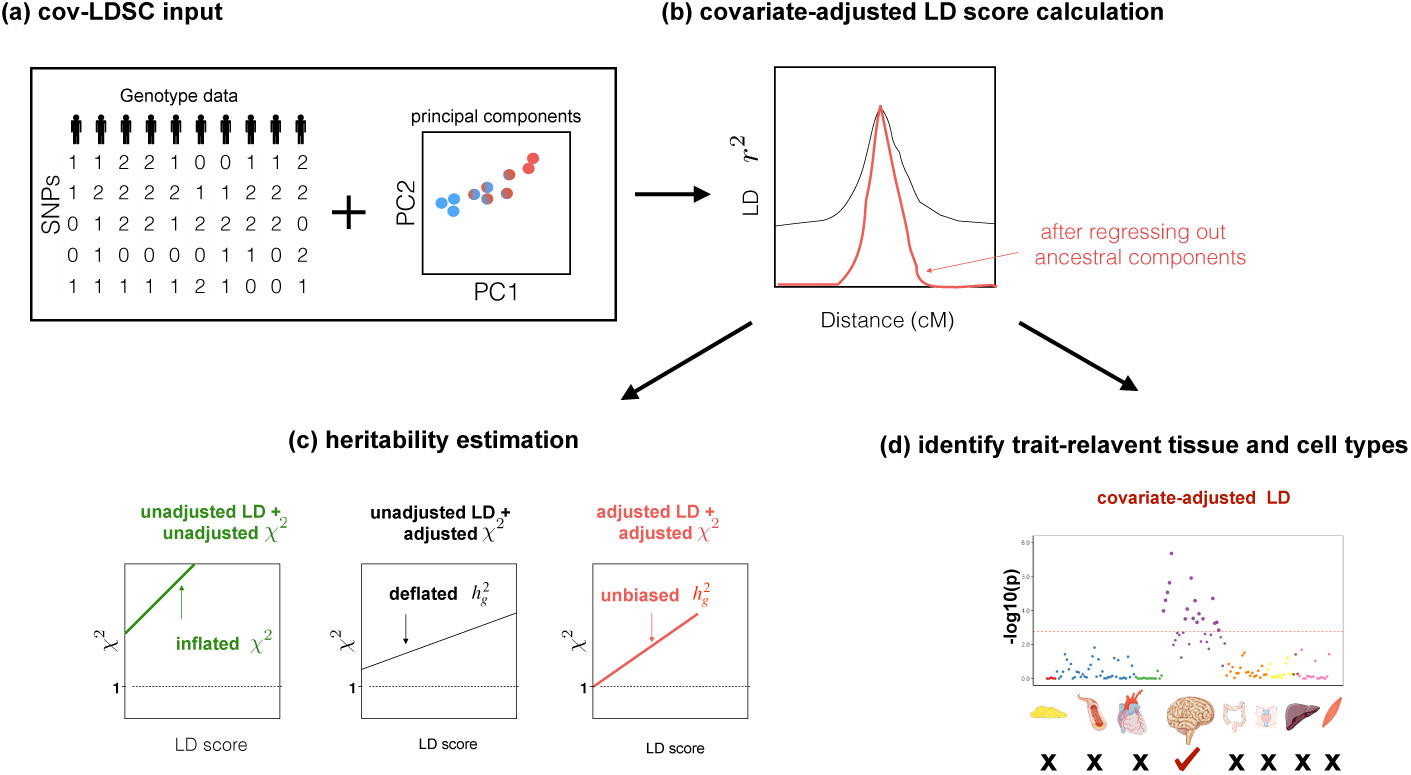
Overview of the covariate-adjusted LD score regression. (a) As input, cov-LDSC takes raw genotypes of collected GWAS samples and their global principal components. (b) cov-LDSC regresses out the ancestral components based on global principal components from the LD score calculation and corrects for long-range admixture LD. Black and red lines indicate estimates before and after covariate adjustment respectively (c) Adjusted heritability estimation based on GWAS association statistics (measured by *χ*^2^) and covariate-adjusted LD scores. (d) Estimation of heritability enrichment in tissue-specific gene sets.

### Robustness of LD score estimation

To demonstrate the effect of admixture on the stability of LD score estimates, we first calculated LD scores with genomic window sizes ranging from 0-50 cM in both European (EUR, *N* = 503) and admixed American (AMR, *N* = 347) populations from the 1000 Genomes Project [20]. As window size increases, we expect the mean LD score to plateau because LD should be negligible for large enough distance. If the mean LD score does not plateau, but continues to rise with increasing window size, then one of two possibilities may apply: (1) the window is too small to capture all of the LD; (2) the LD scores are capturing long-range pairwise SNP correlations arising from admixture. If this increase is non-linear then there is non-negligible distance-dependent LD, violating LDSC assumptions. Examining unadjusted LD scores, we observed that in the EUR population [5], the mean LD score estimates plateaued at windows beyond 1-cM in size, as previously reported. However, in the AMR population the mean LD score estimates continued to increase concavely with increasing window size. In contrast, when we applied cov-LDSC with 10 PCs to calculate covariate adjusted LD scores, we observed that LD score estimates plateaued for both EUR and AMR at a 1-cM and 20-cM window size respectively (< 1% increase per cM, S1 Table). This suggested that cov-LDSC was able to correct the long-range LD due to admixture and yielded stable estimates of LD scores (**Method**, S1 Fig), and also that cov-LDSC was applicable in homogeneous populations (S1 Table). The larger window size for the AMR population was needed due to residual LD caused by recent admixture. We next tested the sensitivity of the LD score estimates with regard to the number of PCs included in the cov-LDSC. We observed that in the AMR panel, LD score estimates were unaffected by adding PCs and by increasing window sizes above 20-cM (S2 Fig).

### Simulations with simulated genotypes

To assess whether cov-LDSC produces less biased estimates of 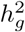, we simulated genotypes of two admixed populations (African American and Latino, **Methods**). We simulated genotypes of 10, 000 unrelated diploid admixed individuals for approximately 400, 000 common SNPs on chromosome 2 in a coalescent framework using msprime [21](**Methods**). First, we tested LDSC and cov-LDSC with different admixture proportions between two ancestral populations, and a quantitative phenotype with a 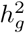 of 0.4 using an additive model (**Methods**). We observed that as the proportion of admixture increased, 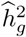 for LDSC increasingly underestimated true 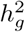 by as much as 18.6%. In marked contrast, cov-LDSC produced consistently less biased estimates regardless of admixture proportion for both Latinos (S3 Fig(a)) and African Americans (S4 Fig). Since both simulated admixed populations would lead to the same conclusions, we performed the subsequent simulations in the Latino individuals only.

Second, we varied the percentage of causal variants from 0.01% to 50% in a polygenic quantitative trait with 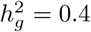 in a population with a fixed admixture proportion of 50%. LDSC again consistently underestimated 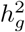 by 12% − 18.6%. In contrast, cov-LDSC yielded less biased estimates regardless of the percentage of causal variants (S3 Fig(b)).

Third, we assessed the robustness of LDSC and cov-LDSC for different assumed total 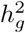 (0.05, 0.1, 0.2, 0.3, 0.4 and 0.5). At each 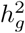 value, LDSC underestimated by 11.5% − 19.6%. For cov-LDSC, we observed that the standard error increased with 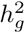, but point estimates remained less biased (S3 Fig(c)).

Fourth, we included an environmental stratification component aligned with the first PC of the genotype data (**Methods**), and concluded that cov-LDSC was also robust to confounding (S3 Fig(d)).

Finally, to assess the performance of cov-LDSC in polygenic binary phenotypes, we simulated genotype data for a binary trait with a prevalence of 0.1 assuming a liability threshold model (**Methods**). We showed that cov-LDSC provided less biased estimates in case-control studies with the same four simulation scenarios (S5 Fig). In contrast, LDSC underestimated heritability for binary phenotypes in the same way as it did for quantitative phenotypes.

### Simulation results with real genotypes

We next examined the performance of both unadjusted LDSC and cov-LDSC on real genotypes of individuals from admixed populations. We used genotype data from the SIGMA cohort, which includes 8,214 Mexican and other Latino individuals. Using ADMIXTURE [22] and populations from the 1000 Genomes Project [20] as reference panels, we observed that each individual in the SIGMA cohort had different admixture proportions (S6 Fig). As in the AMR panel, we observed that using a 20-cM window, LD score estimates plateaued in the SIGMA cohort (S7 Fig, S2 Table), and were unaffected by different numbers of PCs (S8 Fig). When we assumed a non-infinitesimal, additive model with 1% of all SNPs to be causal and 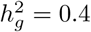, we observed that cov-LDSC 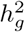 estimates produced less biased estimates using a 20-cM window with 10 PCs (S9 Fig). We subsequently used a 20-cM window and 10 PCs in all simulations.

We observed that cov-LDSC yielded less biased 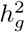 estimates in simulated traits where we varied the number of causal variants and total heritability compared to the original LDSC (**Fig 2(a)-(b)**). In contrast, LDSC underestimated heritability by as much as 62.5%. To examine the performance of cov-LDSC in the presence of environmental confounding factors, we simulated an environmental stratification component aligned with the first PC of the genotype data, representing European v.s. Native American ancestry. In this simulation scenario, cov-LDSC still provided less biased 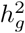 estimates (**Fig 2(c)**). Intercepts of all the simulation scenarios were less than the genomic control inflation factor (GC), suggesting that polygenicity accounts for a majority of the increase in the mean *χ*^2^ statistic compared to potential confounding biases (S10 Fig(a)-(c), S3 Table).

**Fig 2.**
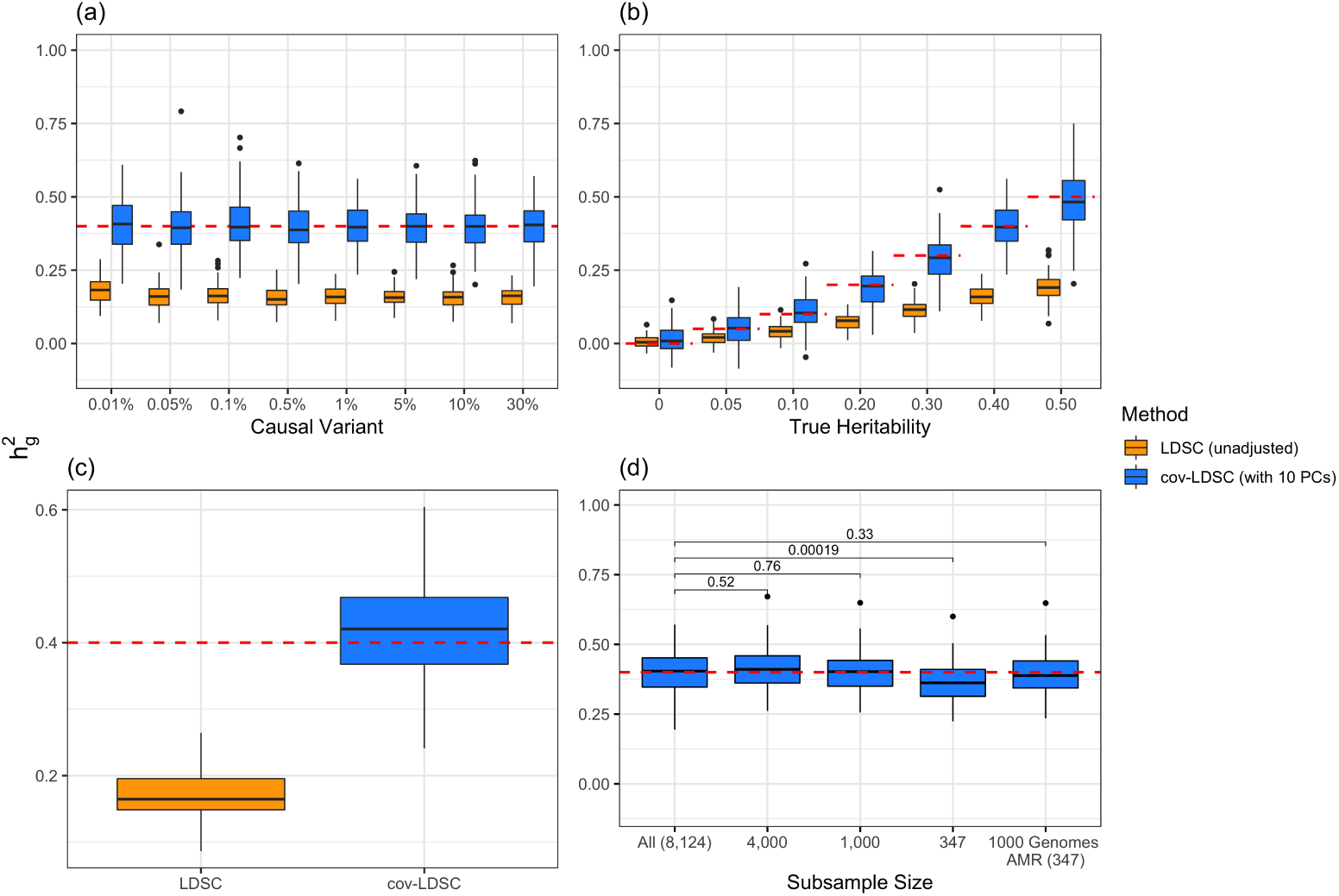
Estimates of heritability 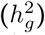 under different simulation scenarios using the SIGMA cohort. LDSC (orange) underestimated 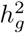 and cov-LDSC (blue) yielded robust 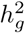 estimates under all settings. Each boxplot represents the mean LD score estimate from 100 simulated phenotypes using the genotypes of 8,214 unrelated individuals from the SIGMA cohort. We used a window size of 20-cM in both LDSC and cov-LDSC, and 10 PCs were included in cov-LDSC in all scenarios. A true polygenic quantitative trait with 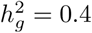 is assumed for scenarios (a), (c) and (d) and 1% causal variants are assumed for scenarios (b)-(d). (a) 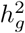 estimation with varying proportions of causal variants (0.01% −30%). (b) 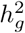 estimation with varying heritabilities (0, 0.05, 0.1, 0.2, 0.3, 0.4 and 0.5). (c) 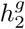 estimation when an environmental stratification component aligned with the first PC of the genotype data was included in the phenotype simulation. (d) 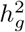 estimation when using a subset of the cohort to obtain LD score estimates and using out-of-sample LD score estimates obtained from Admixed Americans included in the 1000 Genomes Project [20].

Thus far, we have used cov-LDSC by calculating LD scores on the same set of samples that were used for association studies (in-sample LD scores). In practical applications, computing LD scores on the whole data set can be computationally expensive and difficult to obtain, so we investigated computing LD scores on a subset of samples. To investigate the minimum number of samples required to obtain accurate in-sample LD scores, we computed LD scores on subsamples of 100, 500, 1, 000 and 5, 000 individuals from a GWAS of 10, 000 simulated genotypes (S11 Fig). We repeated these analyses in simulated phenotypes in the SIGMA cohort. We subsampled the SIGMA cohort, and obtained less biased estimates when using as few as 1, 000 samples (**Fig 2(d)**). We therefore recommend computing in-sample LD scores on a randomly chosen subset of at least 1, 000 individuals from a GWAS in our approach.

### Assessing power and bias in tissue type specific analysis

Following Finucane et al [9], we extended cov-LDSC so that we can assess enrichment in and around sets of genes that are specifically expressed in tissue and cell-types (cov-LDSC-SEG). To test whether cov-LDSC can produce robust results with properly controlled type I error, We calculated the in-sample LD scores using LDSC and cov-LDSC, respectively, using a 20-cM window and 10 PCs in cov-LDSC for all 53 baseline and limbic system annotations. We used PLINK2 [23] for association test and performed tissue type specific enrichment analysis using both LDSC and cov-LDSC for limbic system conditioning on all 53 baseline annotations. We reported the number of significant tests out of 1, 000 simulations in each scenario. We observed no inflation in false-positive rate (FPR) at 0.05 for both LDSC and cov-LDSC under null (i.e., no enrichment). The greatest gains in power were observed in cases where there were modest enrichment (< 2×). We showed that cov-LDSC-SEG was better powered to detect tissue type specific signals compared to LDSC-SEG (S12 Fig).

### Application to SIGMA and 23andMe cohorts

We next used cov-LDSC to estimate 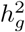 of height, BMI and T2D phenotypes, measured within the SIGMA cohort (**Methods, Table 1**). We estimated 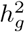 of height, BMI and T2D to be 0.38 ± 0.08, 0.25 ± 0.06 and 0.26 ± 0.07, respectively. These results were similar to reported values from UK Biobank [24] and other studies [17, 25] for European populations. Although estimands differed in different studies (**Methods**), we noted that without cov-LDSC, we would have obtained severely deflated estimates (**Table 1**). To confirm that our reported heritability estimates were robust under different model assumptions, we applied an alternative approach based on REML in the linear mixed model framework implemented in GCTA [17]. To avoid biases introduced from calculating genetic relatedness matrices (GRMs) in admixed individuals, we obtained a GRM based on an admixture-aware relatedness estimation method REAP [26] (**Methods**). GCTA-based results were similar to reported 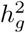 estimates from cov-LDSC, indicating our method was able to provide reliable 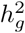 estimates in admixed populations (**Table 1**). We noted, however, that the GCTA-based results would be computationally expensive to obtain on the much larger datasets, for example the 23andMe cohort described below.

**Table 1.** Heritability estimates of height, BMI and type 2 diabetes using different estimation methods. Reported values are estimates of 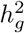 (with standard deviations in brackets) from LDSC using a 20-cM window, cov-LDSC using a 20-cM window and 10 PCs, and GCTA using REAP [26] to obtain the genetic relationship matrix with adjustment by 10 PCs. The final column provides reported 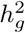 estimates in European populations from various studies [12, 24, 25].

| Phenotype | LDSC<br>(baseline) | cov-LDSC<br>(baseline) | GCTA<br>(REAP) | Public |
| --- | --- | --- | --- | --- |
| **Height** | 0.159 (0.037) | 0.379 (0.079) | 0.450 (0.042) | 0.450-0.685 [12, 24] |
| **BMI** | 0.113 (0.030) | 0.248 (0.061) | 0.235 (0.041) | 0.246-0.270 [24] |
| **T2D** | 0.121 (0.035) | 0.263 (0.073) | 0.376 (0.046) | 0.139-0.414 [24, 25] |

We next applied both LDSC and cov-LDSC to 161, 894 Latino, 46, 844 African American and 134, 999 European research participants from 23andMe. We analyzed three quantitative and four dichotomous phenotypes (**Methods**, S4 Table). In this setting, we noted that if different individuals were included in different traits of interests, one would need to re-compute the GRM for each trait when using genotype-based methods such as GCTA [17] or BOLT-REML [19]. Whereas for cov-LDSC we do not require complete sample overlap between LD reference panel and summary statistics generation. Thus one would only need to compute covariate-adjusted baseline LD score once for each cohort. This makes cov-LDSC a more computationally attractive strategy for estimating heritability and its enrichment in large cohorts. We used a 20-cM window and 10 PCs in LD score calculations for both populations (S13 Fig, S5 Table). LDSC and cov-LDSC produced similar heritability estimates in the European population, whereas in the admixed populations, LDSC consistently provided low estimates of 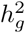 (S6 Table). For each phenotype, we estimated 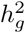 using the same population-specific in-sample LD scores. Intercepts of all the traits were substantially less than the genomic control inflation factor (λ_*gc*_), suggesting that polygenicity accounts for a majority of the increase in the mean *χ*^2^ statistics (S7 Table). For most phenotypes, the reported 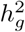 was similar among the three population groups with a notable exception for age at menarche (**Fig 3**, S8 Table). This suggested possible differences (two-sample t-test *p* = 7.1 × 10^−3^ between Latinos and Europeans) in the genetic architecture of these traits between different ancestral groups. It has been long established that there is population variation in the timing of menarche [27–29]. Early menarche might influence the genetic basis of other medically relevant traits since early age at menarche is associated with a variety of chronic diseases such as childhood obesity, coronary heart disease and breast cancer [30, 31]. These results highlighted the importance of including diverse populations in genetic studies in order to enhance our understanding of complex traits that show differences in their genetic heritability.

**Fig 3.**
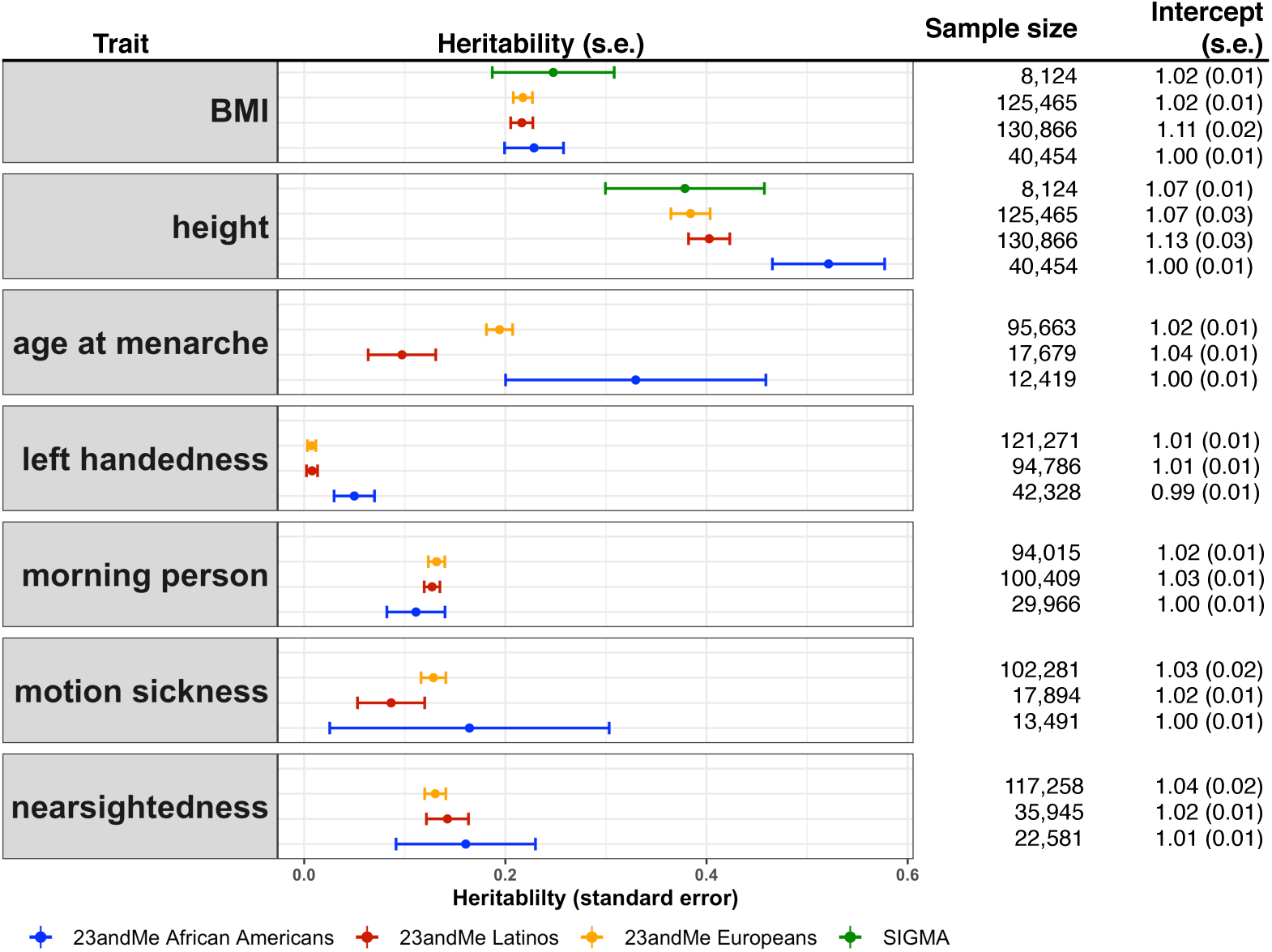
Estimates of heritability 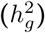 of three quantitative and four dichotomous traits in two admixed populations in the 23andMe research cohort. For seven selected non-disease phenotypes (body mass index (BMI), height, age at menarche, left handedness, morning person, motion sickness and nearsightedness) in the 23andMe cohort, we reported their estimated genetic heritability and intercepts (and their standard errors) using the baseline model. LD scores were calculated using 134, 999, 161, 894, 46, 844 individuals from 23andMe European, Latino and African American individuals respectively. For each trait, we reported the sample size in obtained summary statistics used in cov-LDSC. For BMI and height, we also reported the 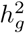 estimates from the SIGMA cohort.

### Tissue type specific analysis

We applied stratified cov-LDSC to sets of specifically expressed genes [9] (SEG) to identify trait-relevant tissue and cell types in traits included in the 23andMe cohort across European, Latino, and African American populations. We only tested height, BMI and morning person, which were the three traits that had heritability z-scores larger than seven in at least two populations [6] (S9 Table). We also performed inverse-variance weighting meta-analysis across the three populations (S10 Table). Across different populations, BMI showed consistent enrichment in central nervous system gene sets. In the European population, most of the enrichments recapitulated the results from the previous analysis using UK Biobank [9]. We found similar but fewer enrichments in Latinos and African Americans, most likely due to smaller sample sizes. The most significantly enriched tissue types for BMI in all three populations were limbic system (*τ*\* _EUR_ = 0.18, *τ*\* _LAT_ = 0.16, *τ*\* _AA_ = 0.28, *τ*\* _meta_ = 0.18), entorhinal cortex (*τ*\* _EUR_ = 0.18, *τ*\* _LAT_ = 0.15, *τ*\* _AA_ = 0.24, *τ*\* _meta_ = 0.17), and cerebral cortex (*τ*\* _EUR_ = 0.16, *τ*\* _LAT_ = 0.14, *τ*\* _AA_ = 0.15, *τ*\* _meta_ = 0.15); none of the three effects were significantly different across populations. When we compared the enrichment for all of the tissues between population pairs, we observed that they have significant non-zero concordance correlation coefficient (*ρ*_EUR-LAT_ = 0.78 (0.72 − 0.83); *ρ*_EUR-LAT_ = 0.32 (0.21 − 0.42)) (**Fig 4(a)-(e)**, S11 Table). The sizes of these three brain structures have been shown to be correlated with BMI using magnetic resonance imaging data [32]. The midbrain and the limbic system are highly involved in the food rewarding signals through dopamine releasing pathway [33]. Furthermore, the hypothalamus in the limbic system releases hormones that regulate appetite, energy homeostasis and metabolisms, like leptin, insulin, and ghrelin [33, 34]. For height, similar to previously reported associations [9], we also identified enrichments in the gene sets derived from musculoskeletal and connective tissues. In the meta-analysis, the three most significant enrichments were cartilage (*τ*\* _EUR_ = 0.21, *τ*\* _LAT_ = 0.19, *τ*\* _AA_ = 0.24, *τ*\* _meta_ = 0.20), chondrocytes (*τ*\* _EUR_ = 0.21, *τ*\* _LAT_ = 0.15, *τ*\* _AA_ = 0.11, *τ*\* _meta_ = 0.17), and uterus (*τ*\* _EUR_ = 0.17, *τ*\* _LAT_ = 0.15, *τ*\* _AA_ = 0.16, *τ*\* _meta_ = 0.16). A heterogeneity test revealed no difference across three populations (*I*^2^ < 70% and p-value > 0.05). The concordance correlation coefficients were *ρ*_EUR-LAT_ = 0.91 (0.89 − 0.93) between European and Latio; *ρ*_EUR-AA_ = 0.60 (0.50 − 0.68) between European and African American (**Fig 4(f)-(j)**, S11 Table). The importance of these tissues and their roles in height have been addressed in the previous pathway analysis, expression quantitative trait loci (eQTLs) and epigenetic profiling [35, 36]. Previous studies have shown that the longitudinal growth of bones is partly controlled by the number and proliferation rate of chondrocytes on the growth plate which is a disc of cartilages [37]. For the morning person phenotype, we found enrichments in many brain tissues in Europeans, concordant with a previous study [38]. Entorhinal cortex (*τ*\* _EUR_ = 0.16, *τ*\* _LAT_ = 0.22, *τ*\* _meta_ = 0.18), cerebral cortex (*τ*\* _EUR_ = 0.15, *τ*\* _LAT_ = 0.22, *τ*\* _meta_ = 0.18), and brain (*τ*\* _EUR_ = 0.17, *τ*\* _LAT_ = 0.19, *τ*\* _meta_ = 0.18) were enriched in both Latinos and Europeans. Evidence showed that circadian rhythm was controlled by the suprachiasmatic nucleus, the master clock in our brain, and also the circadian oscillator that resides in neurons of the cerebral cortex [39–41]. We also found unique enrichments of esophagus muscularis and the esophagus gastroesophageal junction in the Latino populations, but the heterogeneity test showed that the difference is not significant (*I*^2^ = 0.49 and 0.50, respectively). We observed that the concordance correlation coefficient across gene sets was 0.63 (0.51 − 0.68) between Latino and European (**Fig 4(k)-(n)**, S11 Table). Compared to the original LDSC-SEG, cov-LDSC-SEG appeared to have increased statistical power in detecting tissue type specific enrichment in the African American and Latino population (S12 Fig, S14 Fig, S15 Fig, S16 Fig).

**Fig 4.**
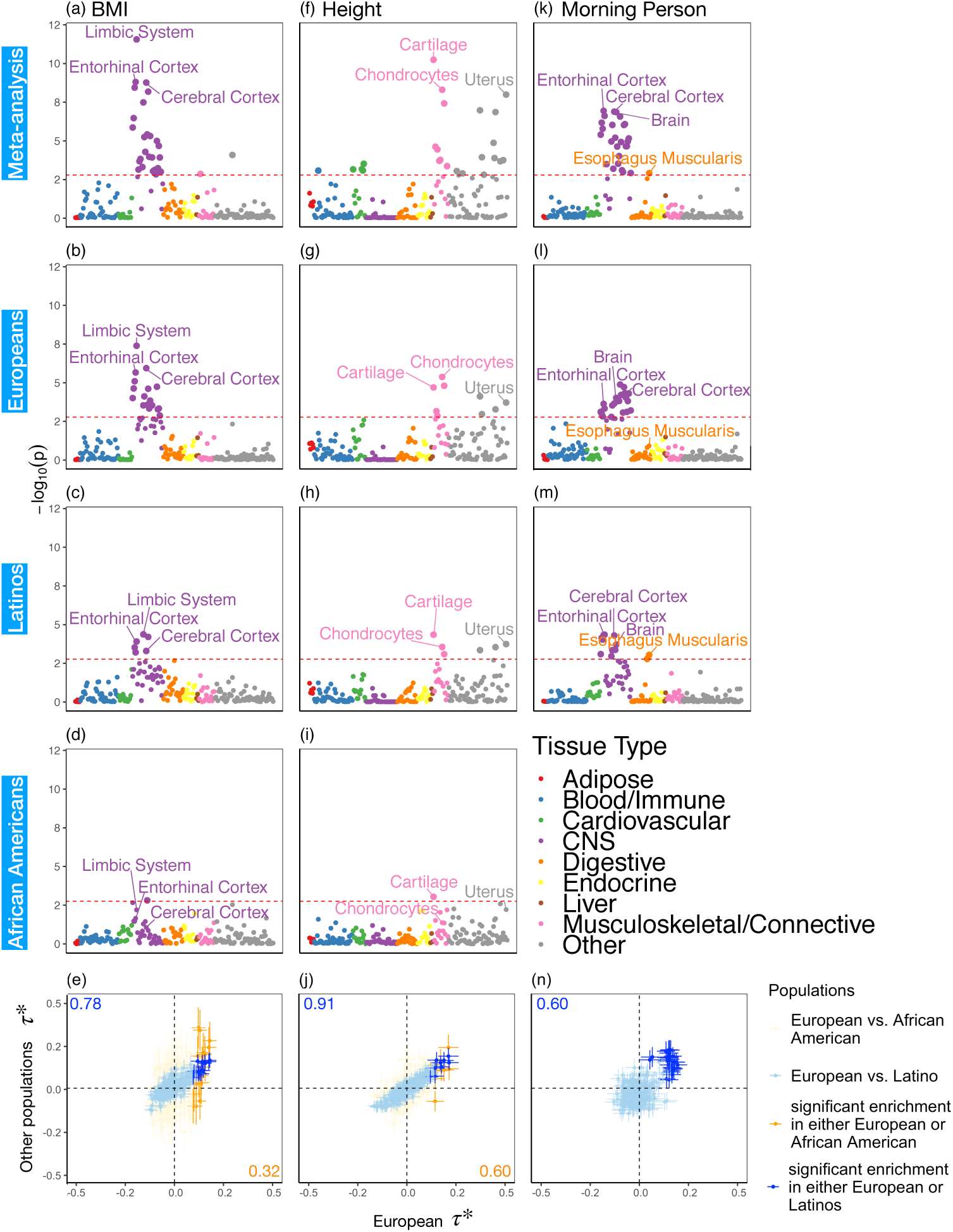
Results of multiple-tissue analysis for height, BMI and morning person. Each point represents a tissue type from either the GTEx data set or the Franke lab data set as defined in Finucane et al [9]. From top to bottom, (a)-(d) show multiple-tissue analysis for BMI in the cross-population meta-analysis and in Europeans, Latinos and African Americans respectively. (e) shows the scatter plot of the estimated per-standardized-annotation effect size *τ*\*, which represents the proportional change of averaged per-SNP heritability for one standard deviation increase in value of the annotation of each cell type, conditional on other 53 non-cell type specific baseline annotations, in the three populations for all tested tissue types (**Methods**). The x-axis shows the *τ*\* in European populations and the y-axis shows either *τ*\* in Latinos (blue) or African Americans (orange). We reported the slope and p-value when we regress Latinos (blue) and African Americans (orange) *τ*\* on Europeans *τ*\* for all tissue types. Error bars indicate standard errors of *τ*\*. Similarly, the results are shown in (f)-(j) for height and (k)-(n) for morning person. The significance threshold in plots (a)-(d), (f-i) and (k-m) is defined by the FDR < 5% cutoff, −log_10_(*p*) = 2.75. Numerical results are reported in S10 Table.

## Discussion

As we expand genetic studies to explore admixed populations around the world, extending statistical genetics methods to make inferences within admixed populations is crucial. This is particularly true for methods based on summary statistics, which are dependent on the use of LD scores that we showed to be problematic in admixed populations. In this study, we confirmed that LDSC that was originally designed for homogenous populations, should not be applied to admixed populations. We introduced cov-LDSC which regresses out global PCs on individual genotypes during the LD score calculation, and showed it can yield less biased LD scores, heritability estimates and its enrichment, such as trait-relevant cell and tissue type enrichments, in homogenous and admixed populations.

Although our work provides a novel, efficient approach to estimate genetic heritability and to identify trait-relevant cell and tissue types using summary statistics in admixed populations, it has a few limitations. First, covariates included in the summary statistics should match the covariates included in the covariate-adjusted LD score calculations (S1 Appendix). To demonstrate this, we simulated the phenotypes using real genotypes included in the SIGMA cohort. We performed cov-LDSC to measure total heritability and its enrichment with varied number of PCs included in summary statistics and in LD score calculation. As the differences between the number of PCs included in the summary statistics and LD score calculation increase, we observed an increase in bias of the total heritability estimation (S17 Fig) and a loss in power when detecting tissue-specific enrichment (S18 Fig). Second, 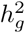 estimates and their enrichment in admixed populations are more sensitive to potentially unmatched LD reference panels. Unmatched reference panels are likely to produce biased estimates [42, 43] and under-powered enrichment analysis (S12 Table, S14 Fig, S15 Fig, S16 Fig). We examined the performance of using an out-of-sample reference panel in admixed populations (See S2 Appendix) and caution that when using 1000 Genomes or any out-of-sample reference panels for a specific admixed cohort, users should ensure that the demographic histories are shared between the reference and the study cohort. Large sequencing projects such as TOPMed [44] that include large numbers (*N* > 1, 000) of admixed samples can potentially serve as out-of-sample LD reference panels, although further investigations are needed to study their properties. We therefore advise to compute in-sample LD scores from the full or a random subset of data (*N* > 1, 000) used to generate the admixed GWAS summary statistics when possible. For tissue and cell type-specific analyses, this means one needs to compute covariate-adjusted LD scores for the genome annotations that were derived from the publicly available gene expression data. We have released open-source software implementing our approach based on all genome annotations derived previously (**URLs**). We strongly encourage cohorts to release their summary statistics and in-sample covariate-adjusted LD scores at the same time to facilitate future studies. Third, when applying cov-LDSC to imputed variants, particularly those with lower imputation accuracy (INFO < 0.99), we caution that the heritability estimates and its enrichment can be influenced by an imperfect imputation reference panel, especially in Latino populations [45, 46]. To limit the bias in varying genotyping array and imputation quality in studied admixed cohorts, we recommend restricting the heritability analyses to common HapMap3 variants. Any extension to a larger set of genetic variants, especially across different cohorts should be performed with caution. Fourth, when we evaluated the performance of cov-LDSC in case-control studies, we assumed no presence of binary covariates with strong effects and demonstrated that cov-LDSC can yield robust 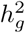 estimates. However, it has been shown that LDSC can provide biased estimates in the presence of extreme ascertainment for dichotomous phenotypes [47]. Adapting cov-LDSC into case-control studies under strong binary effects remains a potential avenue for future work. Fifth, recent studies have shown that heritability estimates can be sensitive to the choice of the LD- and frequency-dependent heritability model [8, 11, 13, 48]. Since our approach can flexibly add annotations to estimate heritability under the model that is best supported by the data, we believe it provides a good foundation for addressing the question of how to incorporate ancestry-dependent frequencies in the LD-dependent annotation in the future (**Methods**). Sixth, summary statistics derived from linear mixed models cannot currently be used for cov-LDSC analysis (S19 Fig). This is due to the fact that, just as the LD needs to be adjusted for the same covariates included in the summary statistics (S1 Appendix), it also needs to be corrected appropriately for the random effect. We leave efficient computation of random effect-adjusted LD score to future work.

Despite these limitations, in comparison with other methods, such as those based on restricted maximum likelihood estimation (REML) [17, 19] with an admixture-aware GRM [26], for estimating 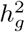 in heterogeneous or admixed populations, cov-LDSC has a number of attractive properties. First, covariate-adjusted in-sample LD scores can be obtained with a subset of samples, enabling analysis of much larger cohorts than was previously possible. Second, LD scores only need to be calculated once per cohort; this is particularly useful in large cohorts such as 23andMe and UK Biobank [49], where multiple phenotypes have been collected per individual and per-trait heritability and its enrichment can be estimated based on the same LD scores. Third, as a generalized form of LDSC, it is robust to population stratification and cryptic relatedness in both homogenous and admixed populations. Fourth, similar to the original LDSC methods, cov-LDSC can be extended to perform analyses such as estimating genetic correlations, partitioning 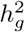 by functional annotations, identifying disease-relevant tissue and cell types and multi-trait analysis [6, 9, 50, 51].

By applying cov-LDSC to approximately 344, 000 individuals from European, African American, and Latin American ancestry, we observed evidence of heritability differences across different populations. Differences in environmental exposures and biological mechanisms can both contribute to the observed differences in genetic heritability across trans-ethnic populations. These differences highlight the importance of studying diverse populations In particular, the differences in biological mechanisms may lead to mechanistic insights about the phenotype. One strategy to do this, which we explored by extending cov-LDSC, is to partition heritability by different cell type- and tissue-specific annotations to dissect the genetic architecture in admixed populations. Our results demonstrated that although there are some cases of nominal heterogeneity across populations among tested tissue-types, most of the tissue-specific enrichments are consistent among the populations studied here. This is consistent with the previous findings that show strong correspondence in functional and cell type enrichment between Europeans and Asians [52, 53]. Seeing the same tissue-type for a single trait emerge in multiple populations can give us more confidence that this tissue may account for polygenic heritability. Larger sample sizes are needed to increase the power of our current analyses and to enhance our understanding of how genetic variants that are responsible for heritable phenotypic variability differ among populations.

As the number of admixed and other diverse GWAS and biobank data become readily available [1, 44, 54], our approach provides a powerful way to study admixed populations.

## Materials and methods

### Mathematical framework of cov-LDSC

Details of the mathematical derivation of cov-LDSC are presented in S1 Appendix. Briefly, in the standard polygenic model on which LDSC is based, *x*_1_, …, *x*_*N*_ are the length-*M* genotype vectors for the *N* individuals, where *M* is the number of SNPs. We model the phenotypes *y*_*i*_

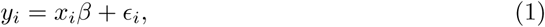

where 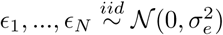 and *β* ∈ ℝ^*M*^ is a vector of per-normalized-genotype effect sizes, which we model as random with mean zero. In standard LDSC, the variance of *β*_*j*_, Var(*β*_*j*_), is the per-SNP heritability of SNP *j*, that is, the total SNP-heritability 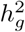 divided by the total number of SNPs *M* (*h*_*g*_^2^*/M*). In stratified LD score regression the variance of *β*_*j*_ depends on a set of genome annotations.

Let *χ*_*j*_^2^ denote the chi-square statistic for the *j*^*th*^ SNP, approximately equal to (*X*_*j*_^*T*^ *Y*)^2^*/N*, where *X*_*j*_ = (*x*_1*j*_, …, *x*_*Nj*_)^*T*^ and *Y* = (*y*_1_, …, *y*_*N*_)^*T*^. The main equation on which LDSC is based is:

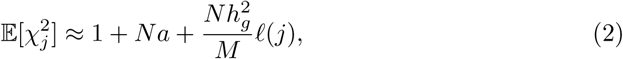

where *α* is a constant that reflects population structure and other sources of confounding, and the LD score, *ℓ* (*j*), is:

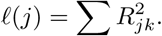

*R*_*jk*_^2^ is the correlation between SNPs *j* and *k* in the underlying population. A new derivation for this equation is given in S1 Appendix. We estimate the total SNP-heritability 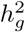 via weighted regression of 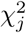 on our estimates of *ℓ* (*j*), evaluating significance with a block jackknife across SNPs [6].

In the absence of covariates, the LD scores can be estimated from an external reference panel such as 1000 Genomes, as long as the correlation structure in the reference panel matches the correlation structure of the sample. In most homogeneous populations, we can also assume that the true underlying correlation is negligible outside of a 1-cM window.

In the presence of covariates, we let *C* denote the *N* × *K* matrix of covariates, each column centered to mean zero, and let *c*_*i*_ be the *i*-th row of *C*. Equation (1) can then be replaced with

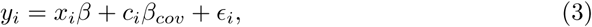

where *β*_*cov*_ is a vector of effect sizes of covariates. We can project the covariates out of this equation by multiplying by *P* = *I* − *C*(*C*^*T*^ *C*)^−1^*C*^*T*^ on the left to get

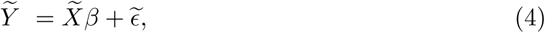

where 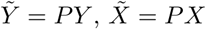 and 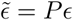 (if the covariates are genotype principal components, then *P* = *I* − *CC*^*T*^).Under this model, an equation identical to Equation (2) can be derived, but where both summary statistics and LD are adjusted for the same covariates (see S1 Appendix).

If *X* is a homogeneous population, then the covariate-adjusted LD will be similar to the non-covariate-adjusted LD and well-approximated by a reference panel. However, if *X* is the genotype matrix from an admixed or heterogeneous population and the covariates include PCs, then the covariate-adjusted LD is no longer well-approximated by either non-covariate-adjusted LD or by a reference panel. Thus, in cov-LDSC, we compute LD scores directly from the covariate-adjusted in-sample genotypes or a random subsample thereof. We call them the covariate-adjusted LD scores.

Using genotype data to compute LD scores means that the model being fit is based on the joint effects of a sparser set of SNPs, e.g. the genotyped SNPs, than when sequence data is used to compute LD scores. For estimating total SNP-heritability, this means that cov-LDSC estimates the same estimand as GCTA 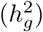 and not the usual estimand of LDSC (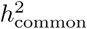; see below). For partitioned heritability, the density of reference panel SNPs can be important because the joint effect of a SNP in an annotation can include the tagged effect of an untyped SNP that is not in the annotation, deflating estimates of enrichment. Thus, we recommend using cov-LDSC only on annotations made of large contiguous regions, such as gene sets. Moreover, we urge caution when interpreting quantitative estimates of heritability enrichment. Here, we look at the significance of the conditional enrichment (i.e., regression coefficient) of gene sets for our tissue-specific analysis (see below).

### Window size and number of PCs in LD score calculations

In addition to computing LD from the covariate-adjusted genotypes, we also investigate the appropriate window size for estimating LD scores. To do this, we examine the effect of varying the genomic window size for both simulated and real data sets. We determine that LD score estimates were robust to the choice of window size if the increase in the mean LD score estimates was less than 1% per cM beyond a given window. Using this criterion, we use window sizes of 5-cM and 20-cM for the simulated and real genotypes, respectively (S13 Table, S2 Table, S5 Table). We also calculate the squared correlations between LD score estimates using the chosen window size and other LD score estimates with window sizes larger than the chosen window. The Pearson squared correlations were greater than 0.99 in all cases (S14 Table, S15 Table, S16 Table) indicating the LD score estimates were robust at the chosen window sizes.

Similarly, to determine the number of PCs needed to be included in the GWAS association tests and cov-LDSC calculations, we examine the effect of varying the genomic window size using different numbers of PCs. The number of PCs that needed to be included for covariate adjustment depended on the population structure for different datasets.

### Genotype simulations

We evaluate the performance of LDSC and cov-LDSC with simulated phenotypes and both simulated and real genotypes. For the simulated genotypes, we used msprime [21] version 0.6.1 to simulate population structure with mutation rate 2 × 10^−8^ and recombination maps from the HapMap Project [55]. We adapt the demographic model from Mexican migration history [56] for Latinos and the out of Africa model [57] for African Americans using parameters that were previously inferred from the 1000 Genomes Project [20]. We assume the admixture event happened approximately 500 years and 200 years ago for Latino and African American populations, respectively. We set different admixture proportions to reflect different admixed populations. In each population, we simulate 10, 000 individuals after removing second degree related samples (kinship> 0.125) using KING [58].

### Slim Initiative in Genomic Medicine for the Americas (SIGMA) Type 2 Diabetes (T2D) cohort

8, 214 Mexican and other Latin American samples were genotyped with Illumina HumanOmni2.5 array. We further filter the genotyped data to be MAF > 5% and remove SNPs in high LD regions. After QC, a total of 8, 214 individuals and 943, 244 SNPs remain. We estimate the in-sample LD score with a 20-cM window and 10 PCs in all scenarios.

We use these genotypes for simulations. We also analyze three phenotypes from the SIGMA cohort: height, BMI, and type 2 diabetes (T2D). For T2D, we assume a reported prevalence in Mexico of 0.144 [16]. For each phenotype, we include age, sex, and the first 10 PCs as fixed effects in the association analyses.

### Phenotype simulations

We simulate phenotypes with two different polygenic genetic architectures, given by GCTA [17] and the baseline model [6], respectively. In the GCTA model, all variants are equally likely to be causal independent of their functional or minor allele frequency (MAF) structure, and the standardized causal effect size variance is constant, i.e. 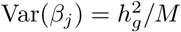. In contrast, the baseline model incorporates functionally dependent architectures. Briefly, it includes 53 overlapping genome-wide functional annotations (e.g. coding, conserved, regulatory). It models Var(*β*_*j*_) = ∑_*C*_ *α*_*c*_(*j*) *τ*_*c*_ where *α*_*c*_(*j*) is the value of annotation *α*_*c*_ at variant *j* and *τ*_*c*_ represents the per-variant contribution, of one unit of the annotation *α*_*c*_, to heritability. We generate all causal variants among common observed variants with MAF > 5% (∼ 40, 000 SNPs in simulated genotypes and 943, 244 SNPs in the SIGMA cohort). To represent environmental stratification, similar to previously described [5], we add 0.2× standardized first principal component to the standardized phenotypes.

We simulate both quantitative and case-control traits with both GCTA and baseline model genetic architectures, using both simulated and real genotypes, varying the number of causal variants, the true heritability, and environmental stratification. For case-control simulations, we adopt a liability threshold model with disease prevalence 0.1. We obtain 5, 000 cases and 5, 000 controls for each simulation scenario.

To obtain summary statistics for the simulated traits, we apply single-variant linear models for quantitative traits and logistic models for binary trait both with 10 PCs as covariates in association analyses using PLINK2 [23].

### 23andMe cohort

All participants were drawn from the customer base of 23andMe, Inc., a direct to consumer genetics company. Participants provided informed consent and participated in the research online, under a protocol approved by the external AAHRPP-accredited IRB, Ethical & Independent Review Services (www.eandireview.com). Samples from 23andMe are then chosen from consented individuals who were genotyped successfully on an Illumina Infinium Global Screening Array (∼ 640, 000 SNPs) supplemented with∼ 50, 000 SNPs of custom content. We restrict participants to those who have European, African American, or Latino ancestry determined through an analysis of local ancestry [59].

To compute LD scores, we use both genotyped and imputed SNPs. We filter genotyped variants with a genotype call rate ≤ 90%, non-zero self-chain score, strong evidence of Hardy Weinberg disequilibrium (*p* > 10^−20^ to accommodate large sample sizes included for detecting deviations), and failing a parent-offspring transmission test. For imputed variants, we use a reference panel that combined the May 2015 release of the 1000 Genomes Phase 3 haplotypes [20] with the UK10K imputation reference panel [60]. Imputed dosages are rounded to the nearest integer (0, 1, 2) for downstream analysis. We filter variants with imputation r-squared ≤ 0.9. We also filter genotyped and imputed variants for batch effects (if an F-test from an ANOVA of the SNP dosages against a factor dividing genotyping date into 20 roughly equal-sized buckets has a p-value less than 10^−50^) and sex dependent effects (if the r-squared of the SNP is greater than 0.01 after fitting a linear regression against the gender). To minimize rounding inaccuracies, we prioritize genotyped SNPs over imputed SNPs in the merged SNP set. We restrict the merged SNP set to HapMap3 variants with MAF ≥ 0.05. We measure LD scores in a subset of African Americans (61, 021) and Latinos (9, 990) on chromosome 2 with different window sizes from 1-cM to 50-cM (S5 Table) and squared correlation between different window sizes (S16 Table). We compute all LD scores with a 20-cM window.

In genome-wide association analyses, for each population, we choose a maximal set of unrelated individuals for each analysis using a segmental identity-by-descent (IBD) estimation algorithm [61]. We define individuals to be related if they share more than 700-cM IBD.

We perform association tests using linear regression model for quantitative traits and logistic regression model for binary traits assuming additive allelic effects. We include covariates for age, sex and the top 10 PCs to account for residual population structure. We list details of phenotypes and genotypes in S4 Table.

### Heritability estimation

We calculate in-sample LD scores using both a non-stratified LD score [5] model and the baseline model [6]. In simulated phenotypes generated with the GCTA model, we use non-stratified LDSC to estimate heritability. In simulated phenotypes generated using the baseline model, we use LDSC-baseline to estimate heritability. We use the 53 non-frequency dependent annotations included in the baseline model to estimate 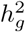 in the 23andMe research database and the SIGMA cohort real phenotypes. We recognize that recent studies have shown that genetic heritability can be sensitive to the choice of LD-dependent heritability model [8, 11, 13]. However, understanding the LD- and MAF-dependence of complex trait genetic architecture is an important but complex endeavor potentially requiring both modeling of local ancestry as well as large sequenced reference panels that are currently unavailable. We thus leave this complexity for future work.

### 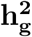 versus 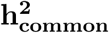

The quantity 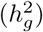 we reported in the main analysis is defined as heritability tagged by HapMap3 variants with MAF ≥ 5%, including tagged causal effects of both low-frequency and common variants. This quantity is different from 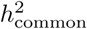, the heritability casually explained by all common SNPs excluding tagged causal effects of low-frequency variants, reported in the original LDSC [5]. In Europeans and other homogeneous populations, it is possible to estimate 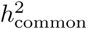, since reference panels, such as 1000 Genomes Project [20], are available which include > 99% of the SNPs with frequency > 1%. However, in-sample sequence data is usually not available for an admixed GWAS cohort, and so cov-LDSC can only include genotyped SNPs in the reference panel, and thus can only estimate the heritability tagged by a given set of genotyped SNPs. In order to compare the same quantity across cohorts, we use common HapMap3 SNPs (MAF ≥ 5%) for in-sample LD reference panel calculation, since most of them should be well imputed for a genome-wide genotyping array. To quantify the difference between 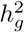 and 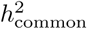, we pre-phase the genotype data in the SIGMA cohort using SHAPEIT2 [62]. We use IMPUTE2 [63] to impute genotypes at untyped genetic variants using the 1000 Genomes Project Phase 3 [20] dataset as a reference panel. We merge genotyped SNPs and all well imputed (INFO> 0.99) SNPs (> 6.9 million) in the SIGMA cohort as a reference panel and reported 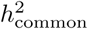, to approximate what the estimate of 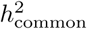 would have been with a sequenced reference panel (S17 Table).

### Tissue type specific analyses

We generate the *τ* for 53 baseline annotations with 40% of annotations with non-zero *τ* and 60% of annotations with zero *τ*. We then generate different regression coefficients *τ* for limbic system in gene sets defined in Franke et al [64, 65] with different enrichment. We scale all the *τ* to make the total 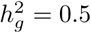. For each variant *j*, the variance of *β*_*j*_ is the sum of the of all the categories that the variant is in (Var(*β*_*j*_) = *τ*_*c*_). We randomly draw *j* from a normal distribution with mean zero and variance 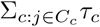 to simulate the phenotypes. We run 1, 000 simulations for each enrichment set (ranging from no (1×) enrichment to 2.5× enrichment). We annotate the genes with the same set of tissue specific expressed genes identified previously [9] using the Genotype–Tissue Expression (GTEx) project [66] and a public dataset made available by the Franke lab [64, 65]. We calculate within-sample stratified cov-LD scores with a 20-cM window and 10 PCs in the 23andMe cohort for each of these 205 gene sets and 53 baseline annotations. We obtain regression coefficients 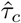 from the model and normalize them as

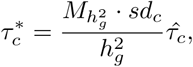

where 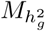 is the number of SNPs used to calculate 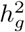 and *sd*_*c*_ is the standard deviation (sd) of annotation *α*_*c*_ [8]. We interpret 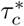 as the proportional change of averaged per-SNP heritability by one sd increase in value of the annotation of each cell type, conditional on other 53 non-cell type specific baseline annotations. We calculate a one-tailed p-value for each coefficient where the null hypothesis is that the coefficient is non-positive [9]. All the significant enrichments are reported with false discovery rate < 5% (− log_10_(*p*) > 2.75). We perform fixed-effect inverse variance weighting meta-analysis using 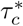 and normalized standard error across populations.

## Software Availability

An open-source software implementation of covariate-adjusted LD score regression is publicly available (see **Web Resources**).

## Web Resources

cov-LDSC software and tutorials, https://github.com/immunogenomics/cov-ldsc

msprime, https://pypi.python.org/pypi/msprime;

GCTA, http://cnsgenomics.com/software/gcta/;

BOLT-LMM, v2.3.4, https://data.broadinstitute.org/alkesgroup/BOLT-LMM/;

LDSC, https://github.com/bulik/ldsc/;

PLINK2, https://www.cog-genomics.org/plink2;

REAP v1.2, http://faculty.washington.edu/tathornt/software/REAP/download.html;

ADMIXTURE v1.3.0,

http://www.genetics.ucla.edu/software/admixture/download.html;

## Acknowledgments

The study was supported by the National Institutes of Health (NIH) TB Research Unit Network, Grant U19 AI111224-01. The content is solely the responsibility of the authors and does not necessarily represent the official views of the NIH.

We thank the research participants of the SIGMA and 23andMe cohort for their contribution to this study.

## Supporting information

**S1 Table. Mean of LD scores with varying window sizes for populations included in the 1000 Genomes project**. AMR (*N* = 347) represents Admixed American and EUR represent European populations (*N* = 503). 10 PCs are included in all cov-LDSC estimates.

**S2 Table. Mean of LD scores with varying window sizes for the SIGMA cohort using LDSC and cov-LDSC**. 10 PCs are included in all cov-LDSC estimates.

**S3 Table. Genomic inflation factor (**λ_*gc*_**), mean chi-square statistics, estimated** 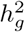 **and intercept under different simulation scenarios using the SIGMA cohort as described in Fig 2 and S10 Fig**. Each estimate represents the mean 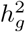 estimates from 100 simulations of 10, 000 unrelated individuals. s.e. represents for standard error.

**S4 Table. Sample sizes (***N* **) and number of SNPs (***M* **) used in LD calculation and heritability estimation of seven selected traits in the 23andMe cohort**.

**S5 Table. mean of LD scores with varying window sizes for the 23andMe cohort using LDSC and cov-LDSC**. 10 PCs are included in all cov-LDSC estimates.

**S6 Table. Heritability estimates of three quantitative and five binary traits included in 23andMe and SIGMA cohorts using different LD models**. Stratified LD model uses genome-wide functional information from all SNPs and explicitly models LD based on 53 functional annotations.

**S7 Table. Heritability estimates, mean chi-square statistics and genomic control inflation factor (**λ_*gc*_**) of three quantitative and four binary traits included in 23andMe using LDSC and cov-LDSC**. cov-LDSC reports the stratified LD model that uses genome-wide functional information from all SNPs and explicitly models LD based on 53 functional annotations.

**S8 Table. Pairwise heritability comparison for seven traits reported in the 23andMe cohort**. P-values are obtained using two-sample t-test with unequal variance.* indicates a p-value passing Bonferroni correction (< 0.05*/*3).

**S9 Table. z-scores for seven traits included in the 23andMe cohort and two continuous traits in the SIGMA cohort**.

**S10 Table. Tissue and type specific analysis on three traits in the 23andMe cohort and their inverse-variance weighting meta-analysis**.

**S11 Table. Concordance correlation coefficient (***ρ***) of pairwise comparison of tissue-type enrichment analysis between two ancestral groups**. We reported the estimated and their 95% confidence intervals (CIs)

**S12 Table. Heritability estimation of seven traits included in the 23andMe Latino cohort when using in-sample and out-of-sample LD reference panel**. We obtain in-sample reference panel from the 23andMe samples and we use 1000 Genomes AMR samples as out-of-sample reference panel. We estimate 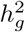 using baseline cov-LDSC model with 10 PCs and a 20-cM.

**S13 Table. Mean of LD scores with varying window sizes for the simulated Latino genotypes using LDSC and cov-LDSC**. 10 PCs are included in all cov-LDSC estimates.

**S14 Table. Pearson r-squared of LD scores with different window sizes when using cov-LDSC in the simulated Latino and African American genotypes**. 10 PCs are included in all cov-LDSC estimates.

**S15 Table. Pearson r-squared of LD scores with different window sizes when using cov-LDSC in the SIGMA cohort**. 10 PCs are included in all cov-LDSC estimates.

**S16 Table. Pearson r-squared of LD scores with different window sizes when using cov-LDSC in the 23andMe cohort**. 10 PCs are included in all cov-LDSC estimates.

**S17 Table. Difference between** 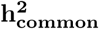 **and** 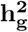 **in the SIGMA cohort for height, body mass index (BMI) and type 2 diabetes (T2D)**.

**S1 Fig. LD score estimates with varying window size in populations from the 1000 Genomes project**. LD score estimates with varying window size using unadjusted LDSC (orange) and cov-LDSC (blue) with 10 PCs with varying window size in both Europeans (*N* = 503, dashed line) and Admixed Americans (*N* = 347, solid line) from the 1000 Genomes Project. The x-axis shows the genomic window size used for estimating LD scores measured in centimorgan (cM). The y-axis shows the mean LD score estimates.

**S2 Fig. LD score estimates with varying window size and number of PCs in Admixed Americans included in the 1000 Genomes project**. LD score estimates (y-axis) using different numbers of PCs at different window sizes (x-axis).

**S3 Fig. Estimates of heritability** 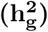 **under different simulation scenarios using the simulated genotypes reflecting a Latino population**. LDSC (orange) underestimated 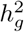 and cov-LDSC (blue) yielded robust estimates under all settings. Each boxplot represents the mean LD score estimate from 100 simulations of 10, 000 unrelated individuals. For cov-LDSC, a window size of 5-cM with 10 PCs are used in all scenarios. For LDSC, a window size of 5-cM are used in all scenarios. A true polygenic quantitative trait with 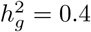 is assumed for scenarios (a), (b) and (d). 1% causal variants are assumed for (a) and (c) - (d). (b)-(d) assumed a dataset with an admixture proportion of 50% from two different ancestral populations. (a) 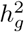 estimation with varying admixed proportions (x-axis) from two ancestral populations. (b) 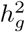 estimation with varying proportions of causal variants (0.01% − 50%). (c) 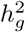 estimation with varying heritability (0.05, 0.1, 0.2, 0.3, 0.4 and 0.5). (d) 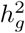 estimation when an environmental stratification component aligned with the first PC of the genotype data is included in the phenotype simulation.

**S4 Fig. Estimates of heritability** 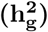 **in simulated genotypes reflecting an African American population**. LDSC (orange) underestimated and cov-LDSC (blue) yielded less biased 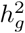 estimates with varying admixed proportions (x-axis). Each boxplot represents the mean LD score estimate from 100 simulations of 10, 000 unrelated African American individuals. For cov-LDSC, a window size of 5-cM with 10 PCs are used in all scenarios. For LDSC, a window size of 5-cM are used in all scenarios. A true polygenic quantitative trait with 1% causal variants and a true 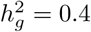 is assumed for scenarios.

**S5 Fig. Estimates of heritability** 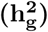 **in case-control phenotypes under different simulation scenarios using the simulated genotypes reflecting a Latino population**. 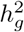 estimation in a phylogenetic binary trait with assumed prevalence of 0.1. 50, 000 unrelated individuals are simulated in total. Each scenario has 5,000 cases and 5,000 controls. 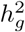 estimation (a) with varying admixed proportions (x-axis) from two ancestral populations; (b) with varying proportions of causal variants (0.01% − 50%); (c) with varying heritability (0.05, 0.1, 0.2, 0.3, 0.4 and 0.5); and (d) when an environmental stratification component aligned with the first PC of the genotype data is included in the phenotype simulation. For cov-LDSC, a window size of 5-cM with 10 PCs are used in all scenarios. For LDSC, a window size of 5-cM are used in all scenarios.

**S6 Fig. ADMIXTURE analysis (K** = **5) of individuals included in the SIGMA cohort and the 1000 Genomes Project**. Each individual is represented as a thin vertical bar. The colors can be interpreted as different ancestries. AFR represents African; AMR represents Admixed American; EAS represents East Asian; EUR represents European and SAS represents South Asian.

**S7 Fig. LD score estimates with varying window size in the SIGMA cohort**. LD score estimates using LDSC (orange) and cov-LDSC (blue) with varying window size in the SIGMA cohort (*N* = 8, 214). The x-axis shows the genomic window size used for estimating LD scores measured in centimorgan (cM). The y-axis shows the mean LD score estimates. For cov-LDSC, 10 PCs are used in all scenarios.

**S8 Fig. LD score estimates with varying window size and number of PCs in the SIGMA cohort**. LD score estimates (y-axis) using different number of PCs at different window sizes (x-axis).

**S9 Fig. Estimates of heritability** 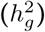 **with varying window sizes used in LD score estimation in the SIGMA cohort**. cov-LDSC (blue) with 10 PCs and varying window size used to obtain LD score. We assumed a true 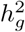 of 0.4 and 1% causal variant in each simulation. 100 replicates are used for each window size.

**S10 Fig. Intercept of estimated** 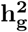 **under different simulation scenarios using the SIGMA cohort as described in Figure 2**. LDSC (orange) underestimated 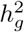 and cov-LDSC (blue) yielded less biased 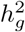 estimates under all settings. Each boxplot represents the mean LD score estimate from 100 simulations of 8, 124 individuals included in the SIGMA project. For cov-LDSC, a window size of 20-cM with 10 PCs are used in all scenarios. For LDSC, a window size of 20-cM are used in all scenarios. A true polygenic quantitative trait with 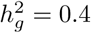 is assumed for scenarios (a), (c) and (d). 1% causal variants are assumed for scenarios (b)-(d). (a) Intercept with varying numbers of causal variants (0.01% − 50%). (b) Intercept with varying heritability (0, 0.05, 0.1, 0.2, 0.3, 0.4 and 0.5). (c) Intercept with the presence of an environmental stratification component aligned with the first PC of the genotype data is included in the phenotype simulation. (d) Intercept when using a subset of total samples and using admixed American samples included in the 1000 Genomes Project.

**S11 Fig. Estimates of heritability** 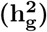 **in simulated genotypes using LD scores estimated with varying sample sizes**. cov-LDSC (blue) is used with varying sample sizes used to obtain LD scores. A random subset of 1%, 5%, 10% and 50% of the total samples (*N* = 10, 000) in the simulated genotypes are used to calculate in-sample LD scores and then to obtain 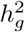 estimates. LD scores are also obtained using independent genotypes (*N* = 1, 000) using the perfect matching demographic model.

**S12 Fig. Simulation results assessing type I error and power for LDSC and cov-LDSC**. We simulate a polygenic trait with 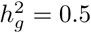. LDSC (orange) shows less power compared to cov-LDSC (blue) in detecting tissue. Each point shows the proportion of simulations (1, 000 for each point) in which a null hypothesis of no tissue enrichment is rejected (Pr(rejected at P ¡ 0.05)), as a function of the z-score of total SNP heritability.

**S13 Fig. LD score estimates with varying window size in populations from 23andMe**. LD score estimates using unadjusted LDSC (orange) and cov-LDSC (blue) with 10 PCs with varying window size in both African Americans (*N* = 46, 844, dashed line) and Latinos (*N* = 161, 894, solid line) from the 23andMe cohort. The x-axis shows the genomic window size used for estimating LD scores measured in centimorgan (cM). The y-axis shows the mean LD score estimates.

**S14 Fig. Tissue and cell type specific analysis with summary statistics in 23andMe Latinos using in-sample original LD and in-sample cov-LD for BMI**. The left panel (a) shows the tissue and cell type specific analysis using original LDSC with in-sample LD scores; while the right panel (b) shows the tissue and cell type specific analysis using cov-LDSC with in-sample cov-LD scores for BMI in 23andMe cohort. The label on the top right in each plot indicates the number of significant tissue type enrichments for each analysis. We observed no difference between LDSC and cov-LDSC in European populations. In contrast, we observed more enrichment in and around sets of genes that are specifically expressed in tissue- and cell-types using cov-LDSC in Latinos and African Americans.

**S15 Fig. Tissue and cell type specific analysis with summary statistics in 23andMe Latinos using in-sample original LD and in-sample cov-LD for height**. The left panel (a) shows the tissue and cell type specific analysis using original LDSC with in-sample LD scores; while the right panel (b) shows the tissue and cell type specific analysis using cov-LDSC with in-sample cov-LD scores for height in 23andMe cohort. The label on the top right in each plot indicates the number of significant tissue type enrichments for each analysis. We observed no difference between LDSC and cov-LDSC in European populations. In contrast, we observed modest increased enrichment using cov-LDSC in Latinos and African Americans.

**S16 Fig. Tissue and cell type specific analysis with summary statistics in 23andMe Latinos using in-sample original LD and in-sample cov-LD for morning person**. The left panel (a) shows the tissue and cell type specific analysis using original LDSC with in-sample LD scores; while the right panel (b) shows the tissue and cell type specific analysis using cov-LDSC with in-sample cov-LD scores for morning person in 23andMe cohort. The label on the top right in each plot indicates the number of significant tissue type enrichments for each analysis. We observed no difference between LDSC and cov-LDSC in European populations. In contrast, we observed modest increased enrichment using cov-LDSC in Latinos and African Americans.

**S17 Fig. Heritability estimate with different number of PCs for GWAS association test and LD score adjustment**. We simulated the phenotypes on the SIGMA cohort using additive model assuming 1% causal SNPs with. We performed univariate cov-LDSC to measure heritability. We varied number of PCs included in summary statistics and varied number of PCs used in cov-LDSC. The x-axis shows the number of PCs included in the cov-LDSC calculation and the y-axis shows the number of PCs included in the summary statistics calculation within the same sample. Numbers in each cell represent the mean estimates from 100 replications. The color (from white to red) represents the statistical difference between the estimated and the truth (measured in − log 10(*P*)). A red cell indicates the 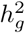 estimate is significantly different from the truth.

**S18 Fig. Type I error in tissue-type-specific enrichment when different number of PCs are used to generate summary statistics and LD scores**. We generated 1, 000 simulations for scenarios where there are different number of PCs (2, 5, 10, 20 and 50) included when calculating LD scores and generating summary statistics (10 PCs) in the cell and tissue-specific enrichment analysis. We simulated a polygenic trait with 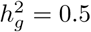. Each bar shows the proportion of simulations in which a null hypothesis of no tissue enrichment is rejected (Pr(rejected at P ¡ 0.05)), as a function of the z-score of total SNP heritability. The horizontal red line indicates *P* = 0.05.

**S19 Fig. LDSC and cov-LDSC with summary statistics derived from linear mixed models**. Estimation of heritability (truth 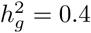) using LDSC and cov-LDSC with 10 (blue) and 50 (green) PCs and a window size of 20-cM. Each boxplot represents the mean LD score estimate from 100 simulations of genotypes from the 8, 124 individuals included in the SIGMA cohort. All summary statistics are derived from linear mixed models with genetic relationship matrix (GRM) only or GRM with 10 genome-wide PCs using GEMMA [67].

**S20 Fig. Results of multiple-tissue analysis for body mass index (BMI), height and type 2 diabetes (T2D) in the SIGMA cohort**. Each point represents a tissue type from either the GTEx data set or the Franke lab data [64, 65]. From left to right, (a)-(d) show multiple-tissue analysis for BMI when using LDSC and cov-LDSC with in-sample and out-of-sample LD reference panels. (e-h) show multiple-tissue analysis for height (e-h) when using LDSC and cov-LDSC with in-sample and out-of-sample LD reference panels. (i-l) show multiple-tissue analysis for T2D when using LDSC and cov-LDSC with in-sample and out-of-sample LD reference panels.

**S21 Fig. Enrichment analysis using in-sample and out-of-sample LD reference panel**. We simulated a polygenic trait with 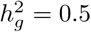. Similar power was obtained when using in-sample (obtained from the SIGMA cohort, turquoise) and out-of-sample (obtained from 1000 Genomes Admixed American (AMR) samples, red) reference panel. In both cases, type I error (at no (1x) enrichment) are well controlled. Each bar shows the proportion of simulations (1,000 for each point) in which a null hypothesis of no tissue enrichment is rejected (Pr(rejected at P ¡ 0.05)), as a function of the z-score of total SNP heritability.

**S22 Fig. Principal component analysis (PCA) of the SIGMA samples**. Samples included in the SIGMA cohort projected onto the first two principal components using SNP weights precomputed from samples in the 1000 Genomes Phase 3 project using SNP weights. AFR represents Africans (green); AMR represents Admixed Americans (orange); EAS represents East Asians (yellow); EUR represents Europeans (blue); SAS represents South Asians (pink) and SIGMA samples are presented in gray.

**S23 Fig. Tissue and cell type specific analysis with summary statistics in 23andMe Latinos using in-sample cov-LD and out-of-sample cov-LD obtained using 1000G AMR samples**. In sample LD is obtained in 23andMe Latinos with 20-cM window size and 10PCs. We observed cell type enrichments in both BMI and height using in-sample cov-LD. However, when we used out of sample 1000G AMR cov-LD with 20cM window size and 10PCs, we observed no cell type enrichments in either BMI and height.

**S24 Fig. Principal component analysis (PCA) of the 23andMe samples**. Samples included in the 23andMe cohort projected onto the first two principal components using SNP weights precomputed from samples in the 1000 Genomes Phase 3 project using SNPweights. AFR represents Africans (green); AMR represents Admixed Americans (red); EAS represents East Asians; EUR represents Europeans (blue); SAS represents South Asians (brown) and the 23andMe samples are presented in gray.

## S1 Appendix Mathematical framework of cov-LDSC

Here, we will first provide a derivation of standard LD score regression that differs somewhat from published derivations, and in particular gives a mathematical interpretation for the value of the intercept. Then we will extend this derivation to cov-LDSC.

### S.1 Review of LD score regression without covariates

#### S.1.1 Summary statistics without covariates

We begin by describing the input data to LD score regresion, which is the output of a standard GWAS.

In a standard GWAS of a quantitative trait, a marginal linear model is fit for each SNP *j*. Let *Y* denote the *N* × 1 vector of phenotypes and *X*_*j*_ denote the *N* × 1 vector of genotypes for SNP *j*, centered to mean zero. In the absence of covariates, we typically fit the model

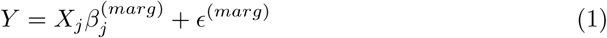

where 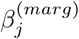 is the marginal effect size of SNP *j* and 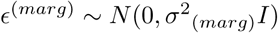.

The F-statistic, which at GWAS sample sizes is approximately chi-square distributed under the null and often referred to as the chi-square statistic, is equal to

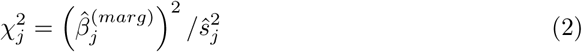

where

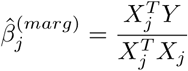

and

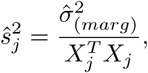

where 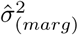 is an estimate of *σ*^2^_(*marg*)_ that, if 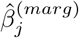 is small, satisfies

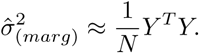

We will assume that 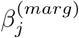 and its estimate 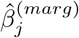 are indeed small, so that this is a valid approximation.

Let 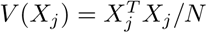 and *V* (*Y*) = *Y*^*T*^ *Y/N* be the empirical variances of *X*_*j*_ and *Y*, and let 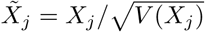, and 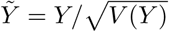 be *X*_*j*_ and *Y*, normalized to empirical variance one. Note that when *X*_*j*_ and *Y* are random, so are 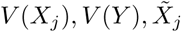, and 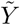. Note also that 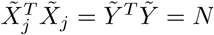, deterministically. We can now simplify the expression for 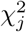:

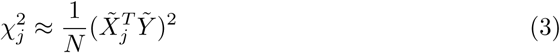

We will assume that we have as input 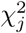 for a genome-wide set of SNPs *j*.

#### S.1.2 The polygenic model

In LD score regression, we take these chi-square statistics as input, and we derive their expectation under a standard polygenic model. Specifically, instead of the marginal model used in GWAS, LD score regression is based on a joint model with random SNP effect sizes:

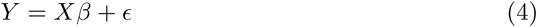

where *Y* is the phenotype vector, *X* = (*X*_1_ … *X*_*M*_) is the *N* × *M* genotype matrix, 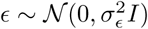, and *β* is the *M* × 1 vector of joint effect sizes. Let 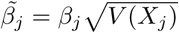, and note that 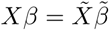. We will model 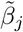 as random with mean zero, independent of each other and of ϵ. Here, we will perform derivations in which 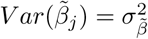; these derivations extend easily to the case in which 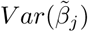 depends on functional annotations. We don’t specify a distribution for 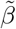.

In LD score regression, we derive the expectation of 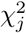 under this polygenic model, and we use the resulting equation to estimate parameters such as 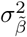. Because *X* is not observed, we ultimately treat it as random. Here, we will derive 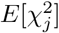 by first deriving 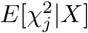 and then using the law of total expectation to remove the conditioning on *X*.

#### S.1.3 Deriving the expression for 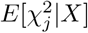

Before deriving the expression for 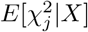, we will first derive the expected empirical variance of *Y*, where the variance is over the random individuals in our GWAS and the expectation is over random *β* and *ϵ*, conditional on *X*.

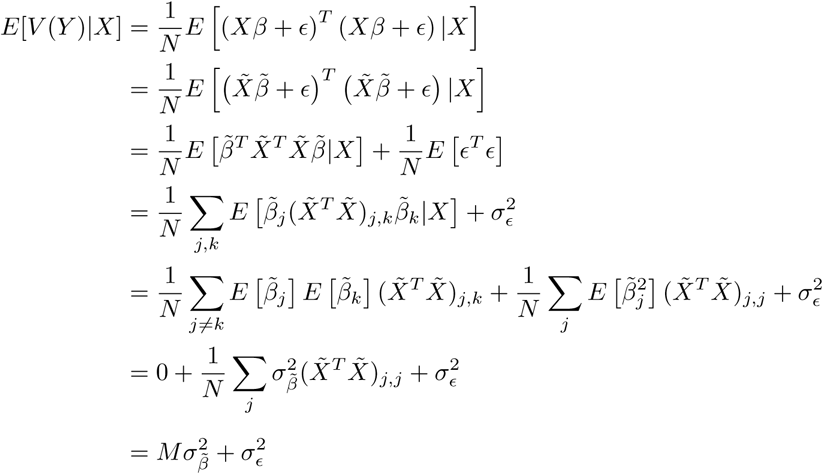

We will let 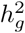 denote 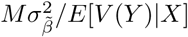, noting that definitions of heritability depend on the model on which they are based, and so 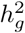 as used here is a different value than in a model in which *β* is fixed.

It will also be useful to have

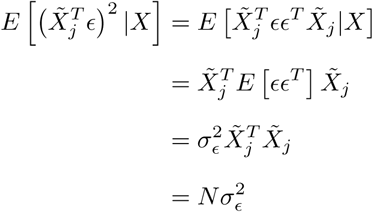

We can now derive the expected chi-square statistic:

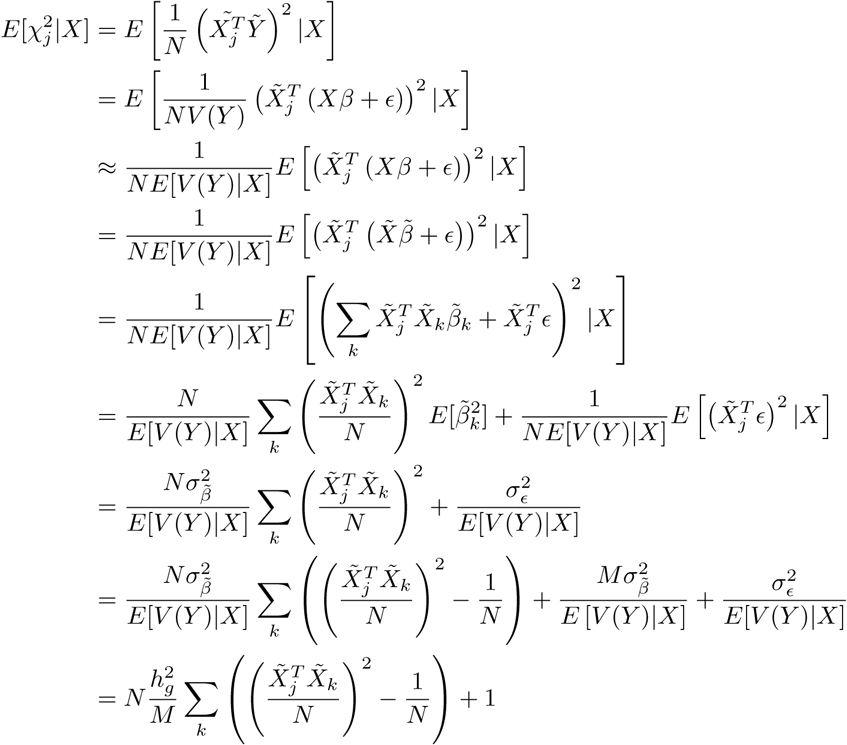

#### S.1.4 Removing the conditioning on *X*

When analyzing summary statistics, we do not have access to the true value of *X*, and so we need to compute the expectation of 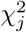 treating *X* as random and integrating it out. To do this, we use the law of total expectation, and so the relevant quantity is 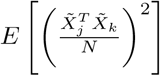. We would like our method to be applicable in the most general circumstances, and so we do not want to assume a particular distribution on *X*, or even that its rows are drawn i.i.d. from some distribution. Instead, we will let *W*_*j*_ denote the set of SNPs in an LD window around *j*, and we will make three assumptions that will allow us to complete our derivation:

1. There is a *c* such that for *k* ∉ *W*_*j*_, we have 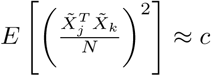, and the approximation is good enough that 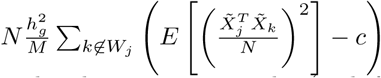 is negligible. If there is no structure or relatedness in our samples (and if *N* is high enough that the difference between standardization in the population and in our sample is negligible), then *c* can be shown to be 1*/N*.
2. For *k* ∈ *W*_*j*_, there is a value *R*_*jk*_ satisfying 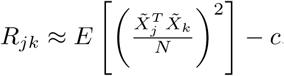, where the approximation is good enough that 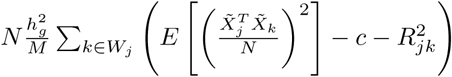 is negligible. Note that if the rows of *X* are drawn i.i.d. from some distribution and *R*_*jk*_ is the correlation between SNPs *j* and *k* in this underlying distribution, and if |*W*_*j*_| is small compared to *M*, then this condition in satisfied.

We can now apply the l o total expec tion t complete the derivation:

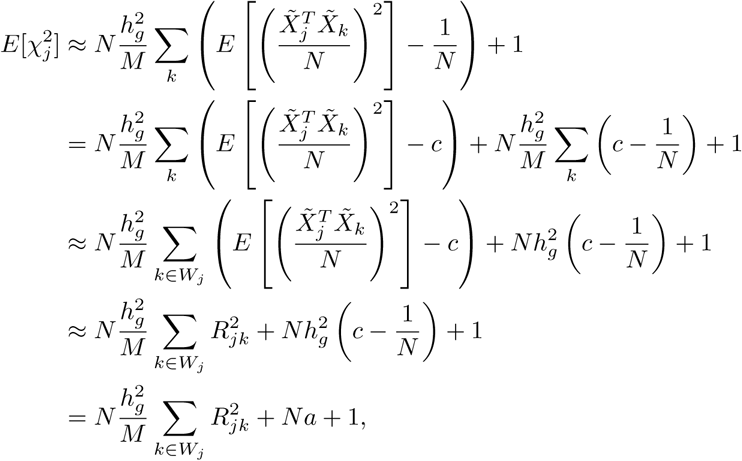

where 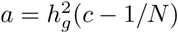. Letting

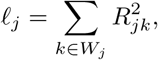

denote the LD score of SNP *j*, we obtain the main LD score regression equation:

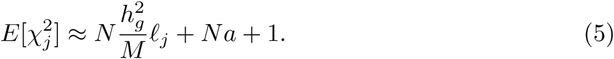

We typically estimate *ℓ*_*j*_ using a reference panel, and we estimate 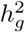 via weighted regression of 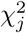 on *ℓ* (*j*), evaluating significance with block jackknife across SNPs.

### S.2 LD score regression in the presence of covariates

We will now discuss LD score regression for a quantitative trait, in the presence of covariates. For a treatment of LD score regression for case-control traits with covariates, see [Weissbrod et al. 2018 AJHG].

#### S.2.1 Summary statistics

Let *C* denote an *N* × *K* matrix of covariates, each column centered to mean zero. In a GWAS of a quantitative trait with covariates, we typically fit the model

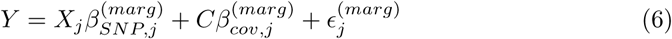

where 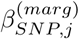 is the marginal effect size of SNP *j* and 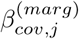 is the effect size vector of the covariates.

The chi-square statistic is equal to

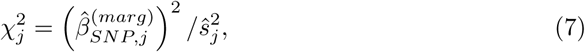

where 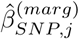 is the least-squares estimate of 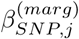, and

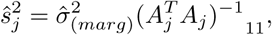

where *A*_*j*_ is the design matrix, given by *A*_*j*_ = (*X*_*j*_ *C*), where 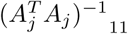 denotes the upper left entry of the matrix 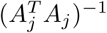, and where 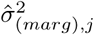 is again an estimate of 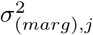.

Let *P* = *I* − *C*(*C*^*T*^ *C*)^−1^*C*^*T*^. By the Frisch-Waugh-Lovell theorem, we have

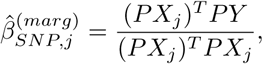

and by block matrix inversion, we have

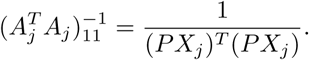

Again assuming that the effect size 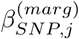 is small, we have

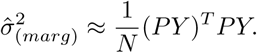

Let *V* (*PX*_*j*_) = ((*P X*_*j*_)^*T*^ *PX*_*j*_)*/N* and *V* (*PY*) = (*PY*)^*T*^ *PY/N*, and let 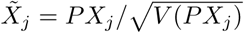, and 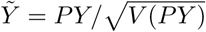. Then, we can rewrite:

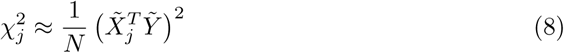

#### S.2.2 Deriving the expression for 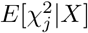

In cov-LDSC, we assume that there are covariates in our GWAS model (Eq (1)) and we include the same set of covariates in the polygenic model that we would like to fit:

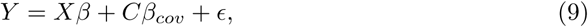

where *Y, X, β, C*, and *ϵ* are as before. Note that under this polygenic model,

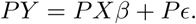

Let 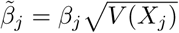. Note that 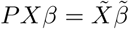.We will model 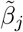 as random with mean zero and variance 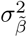. Now we have

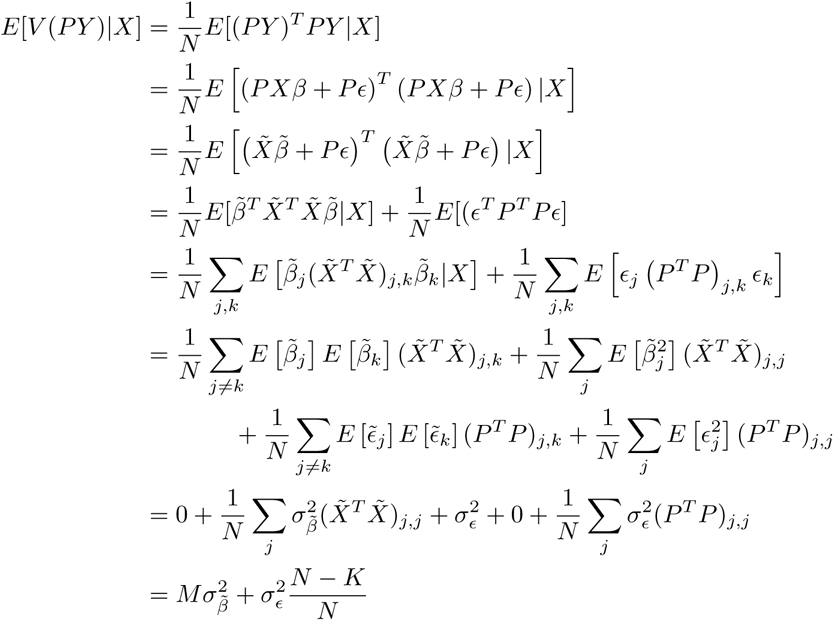

where *K* is the rank of *C*. If *K* is small compared to *N*, as is typical of most GWAS, then we can say that

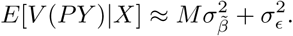

We will let 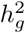 denote 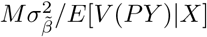. It will again be convenient to have

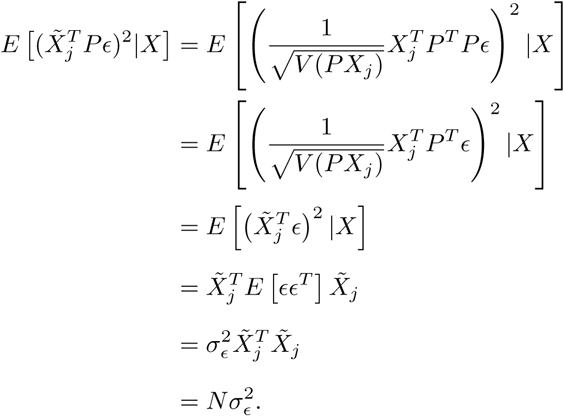

Now we have:

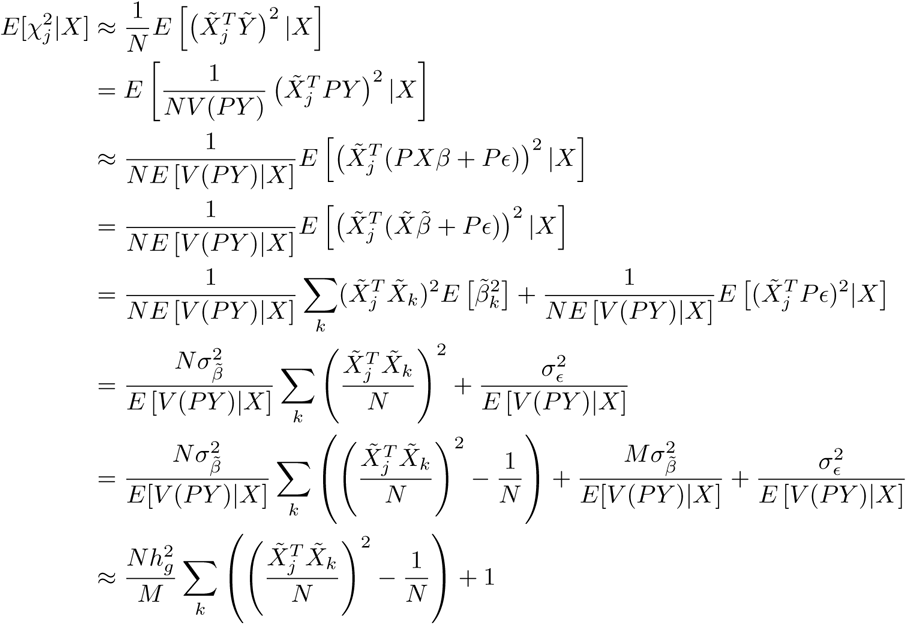

#### S.2.3 Removing the conditioning on *X*

We will make the same two assumptions as for LD score regression without covariates.

1. There is a *c* such that for *k* ∉ *W*_*j*_, we have 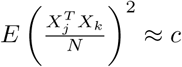. One way to formalize the notion that *C* captures all structure in *X* is that *c* = 1*/N* in this case.
2. For *k* ∈ *W*_*j*_, we have access, for example from a reference panel, to an estimate *R*_*jk*_ satisfying 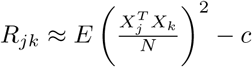. When *X* contains admixture or other structure, correlation as estimated from a reference panel may not suffice. In that case, we can set *R*_*jk*_ to be 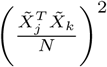, or an estimate of that quantity from a random subsample of the GWAS. We note also that even if window size is 30 cM, this is still only approximately 1% of the genome, and so |*W*_*j*_| is still small compared to *M*.

With these assumptions satisfied, the rest of the derivation is identical to the case without covariates.

## S2 Appendix In-sample versus out-of-sample LD

To test the reliability of using an out-of-sample reference LD panel for cov-LDSC applications, we first examined the performance of out-of-sample LD scores obtained from 1,000 samples with a perfectly matching demographic history in the simulated genotypes. cov-LDSC yielded less biased estimates when using 1,000 samples in an out-of-sample reference panel with a perfectly matching population structure (S11 Fig). Next, we tested the accuracy of heritability estimates and type I error of enrichment analysis when using 1000 Genomes Project [20] Admixed American (AMR) samples to obtain out-of-sample LD scores. When using the AMR panel as a reference panel for the SIGMA cohort, we observed a less biased 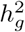 estimate (*P* = 0.33, **Fig 2(d)**), However, as we decreased the number of samples included in the subsampling, the cov-LDSC regression intercepts deviated further from one (S10 Fig(d)). This is probably due to attenuation bias from noisily estimated LD scores at *N* < 1, 000. We observed similar tissue type specific enrichment results for BMI, height and T2D (S20 Fig). We further assessed the power and biases of using 1000 Genomes AMR samples as an external reference panel when applying it in the SIGMA cohort for tissue type specific analysis via simulation. We observed well calibrated type I error and similar power compared to in-sample LD reference panel (S21 Fig). This suggested that the AMR panel included in the 1000 Genomes Project has similar demographic history compared to the SIGMA cohort (S6 Fig, S22 Fig).

Next, we explored the feasibility of applying 1000 Genomes AMR samples in heritability estimation and its enrichment analyses in the 23andMe cohort. We obtained stratified LD scores using 1000 Genomes AMR samples (*N* = 347) and applied it on summary statistics obtained from 23andMe. In contrast to the SIGMA cohort, we discovered total heritability estimates are significantly different from those estimated using in-sample LD scores (S12 Table) and discovered no significant tissue type enrichment (S23 Fig). This suggested that 1000 Genome AMR samples might have different demographic history compared to 23andMe samples (S24 Fig).

We therefore caution that when using 1000 Genomes or any out-of-sample reference panels for a specific admixed cohort, users should ensure that the demographic histories are shared between the reference and the study cohort. We highly recommend computing in-sample LD scores on a randomly chosen subset of at least 1,000 individuals from a GWAS. We also strongly encourage cohorts to release their summary statistics and in-sample covariate-adjusted LD scores at the same time to facilitate future studies.

