## Supplementary_Figures for "Estimating heritability and its enrichment in tissue-specific gene sets in admixed populations"

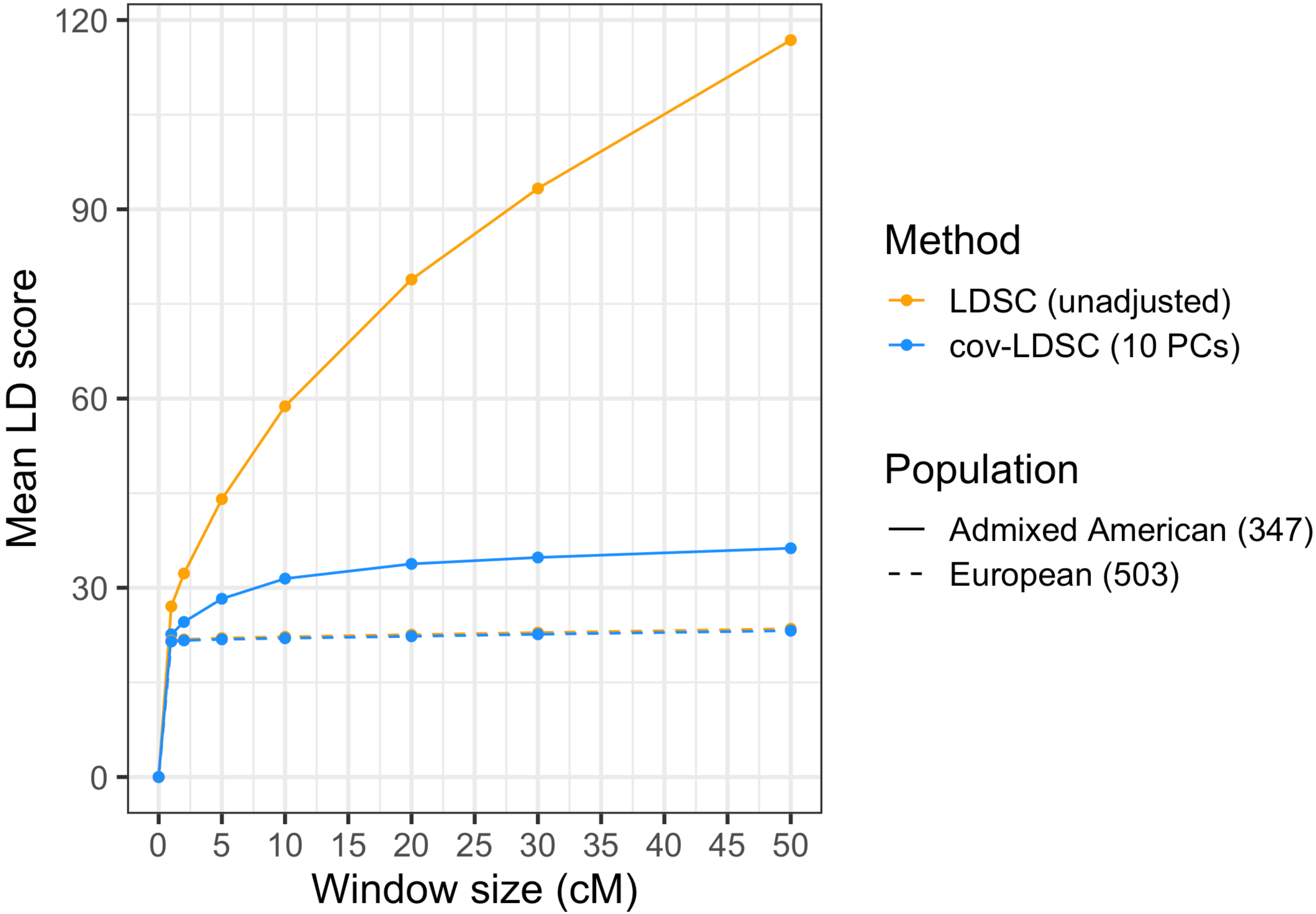


**S1 Fig. LD score estimates with varying window size in populations from the 1000 Genomes project.** LD score estimates with varying window size using unadjusted LDSC (orange) and cov-LDSC (blue) with 10 PCs with varying window size in both Europeans (N=503, dashed line) and admixed Americans (N=347, solid line) from the 1000 Genomes Project. The x-axis shows the genomic window size used for estimating LD scores measured in centimorgan (cM). The y-axis shows the mean LD score estimates.

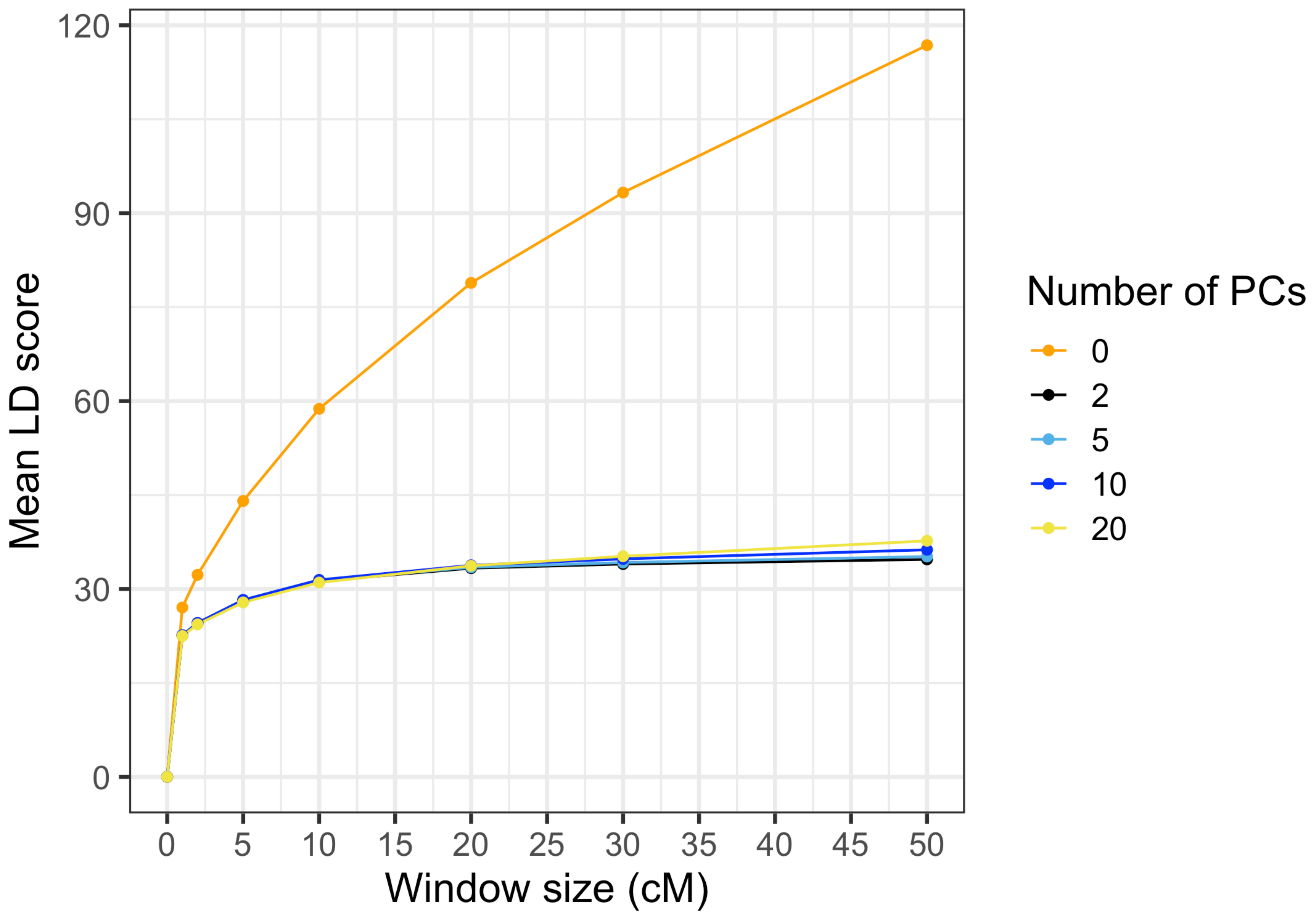


**S2 Fig. LD score estimates with varying window size and number of PCs in admixed Americans included in the 1000 Genomes project.** LD score estimates (y-axis) using different numbers of PCs at different window sizes (x-axis).

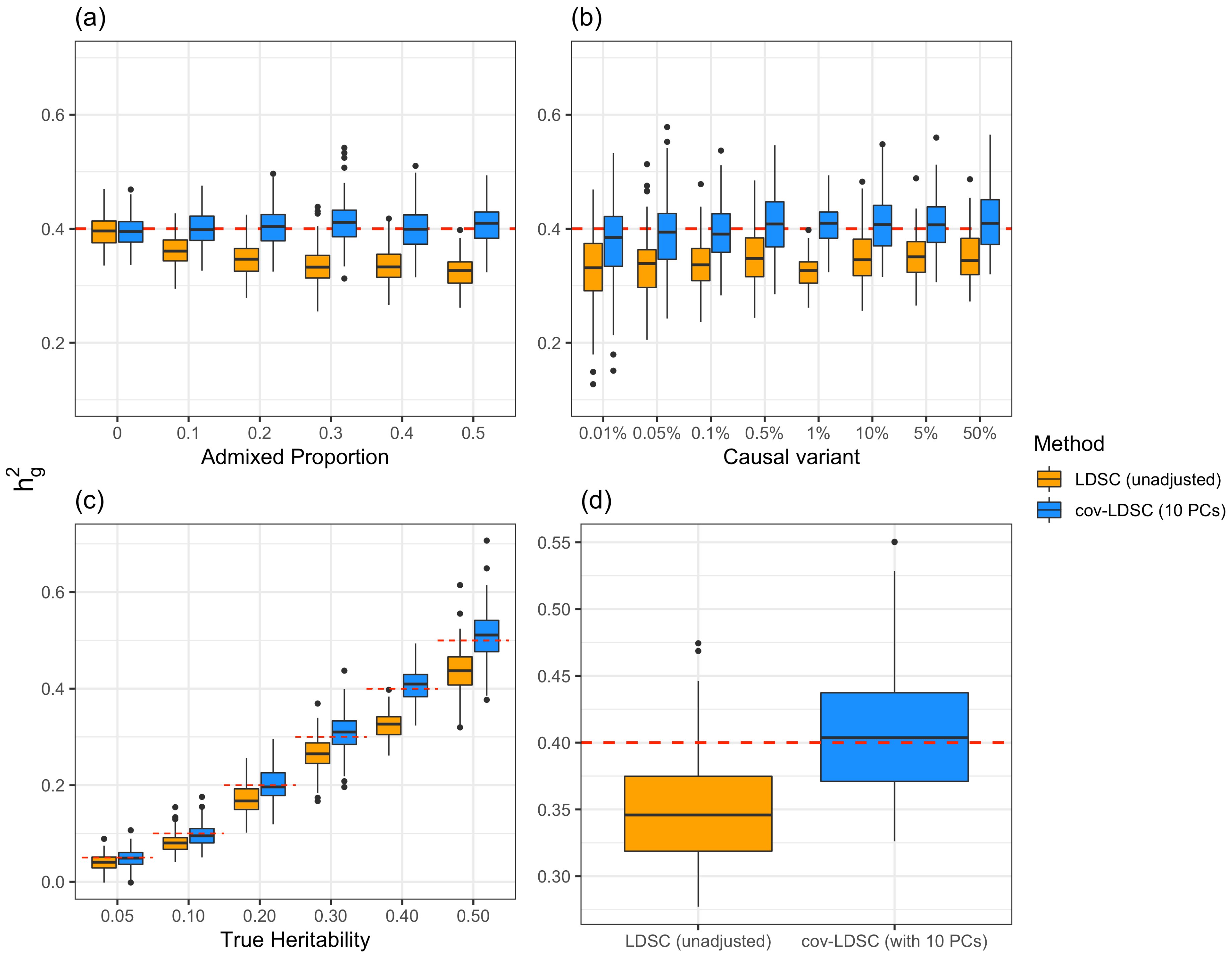


**S3 Fig. Estimates of heritability ($h_g^2$) under different simulation scenarios using the simulated genotypes reflecting a Latino population.** LDSC (orange) underestimated $h_g^2$ and cov-LDSC (blue) yielded robust $h_g^2$ estimates under all settings. Each boxplot represents the mean LD score estimate from 100 simulations of 10,000 unrelated individuals. For cov-LDSC, a window size of 5-cM with 10 PCs are used in all scenarios. For LDSC, a window size of 5-cM are used in all scenarios. A true polygenic quantitative trait with $h_g^2$=0.4 is assumed for scenarios (a), (b) and (d). 1% causal variants are assumed for (a) and (c) - (d). (b)-(d) assumed a dataset with an admixture proportion of 50% from two different ancestral populations. (a) $h_g^2$ estimation with varying admixed proportions (x-axis) from two ancestral populations. (b) $h_g^2$ estimation with varying proportions of causal variants (0.01% - 50%). (c) $h_g^2$ estimation with varying heritabilities (0.05, 0.1, 0.2, 0.3, 0.4 and 0.5). (d) $h_g^2$ estimation when an environmental stratification component aligned with the first PC of the genotype data is included in the phenotype simulation.

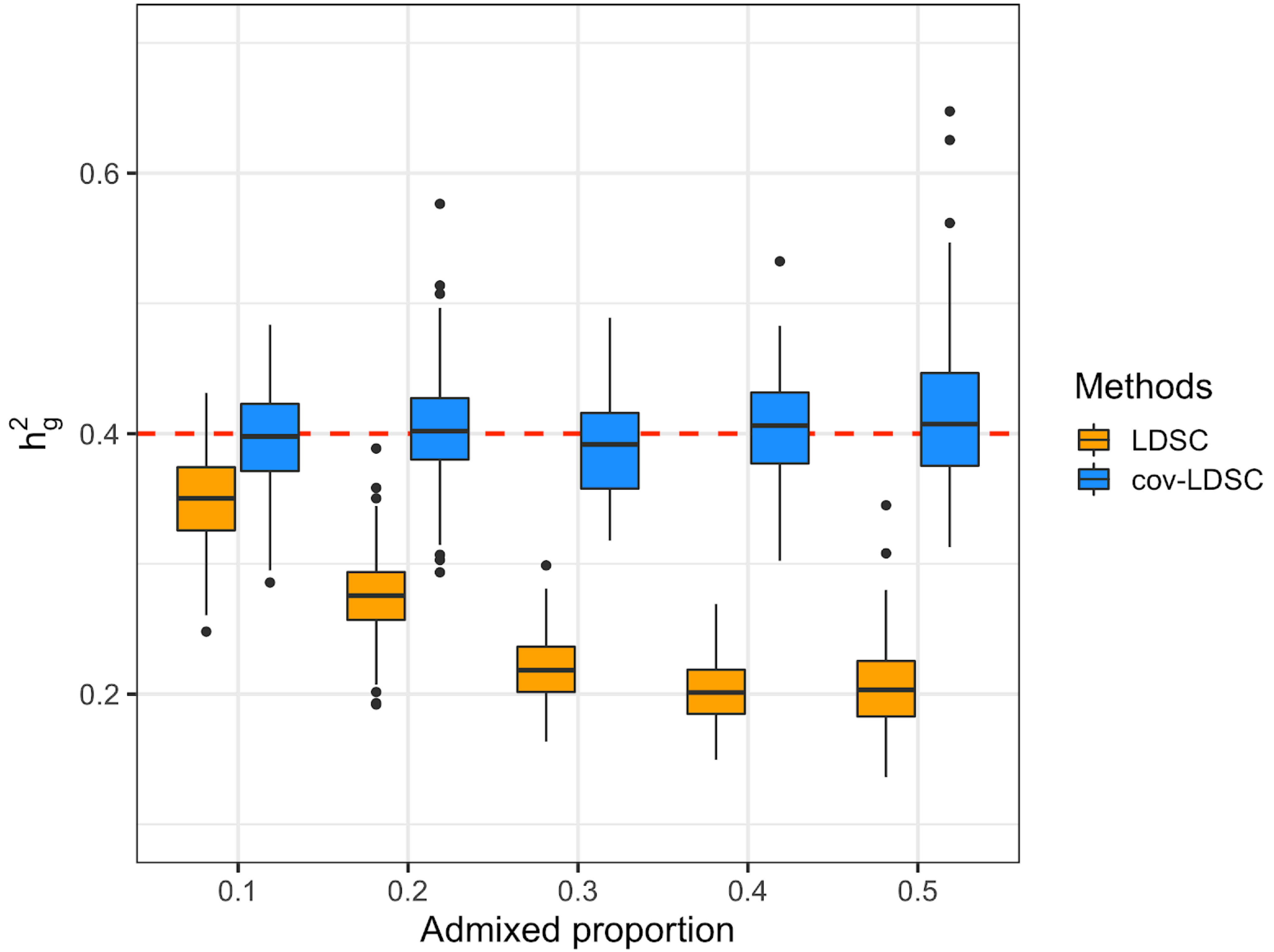


**S4 Fig. Estimates of heritability ($h_g^2$) in simulated genotypes reflecting an African American population.** LDSC (orange) underestimated $h_g^2$ and cov-LDSC (blue) yielded less biased hg2estimates with varying admixed proportions (x-axis). Each boxplot represents the mean LD score estimate from 100 simulations of 10,000 unrelated African American individuals. For cov-LDSC, a window size of 5-cM with 10 PCs are used in all scenarios. For LDSC, a window size of 5-cM are used in all scenarios. A true polygenic quantitative trait with 1% causal variants and a true $h_g^2$=0.4 is assumed for scenarios.

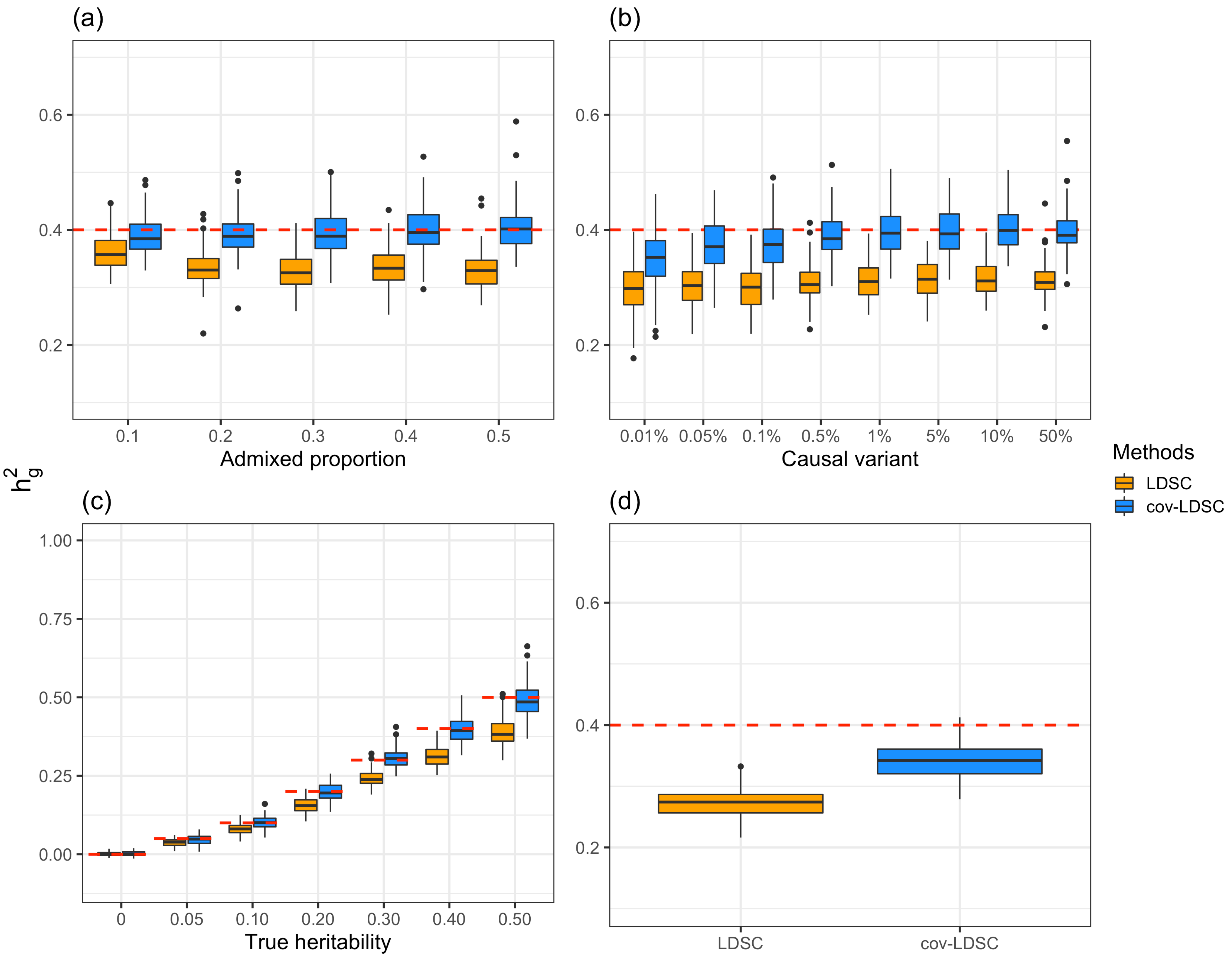


**S5 Fig. Estimates of heritability ( $h_g^2$ ) in case-control phenotypes under different simulation scenarios using the simulated genotypes reflecting a Latino population.** $h_g^2$ estimation in a phylogenetic binary trait with assumed prevalence of 0.1. 50,000 unrelated individuals are simulated in total. Each scenario has 5,000 cases and 5,000 controls. $h_g^2$ estimation (a) with varying admixed proportions (x-axis) from two ancestral populations; (b) with varying proportions of causal variants (0.01% - 50%); (c) with varying heritability (0.05, 0.1, 0.2, 0.3, 0.4 and 0.5); and (d) when an environmental stratification component aligned with the first PC of the genotype data is included in the phenotype simulation. For cov-LDSC, a window size of 5-cM with 10 PCs are used in all scenarios. For LDSC, a window size of 5-cM are used in all scenarios.

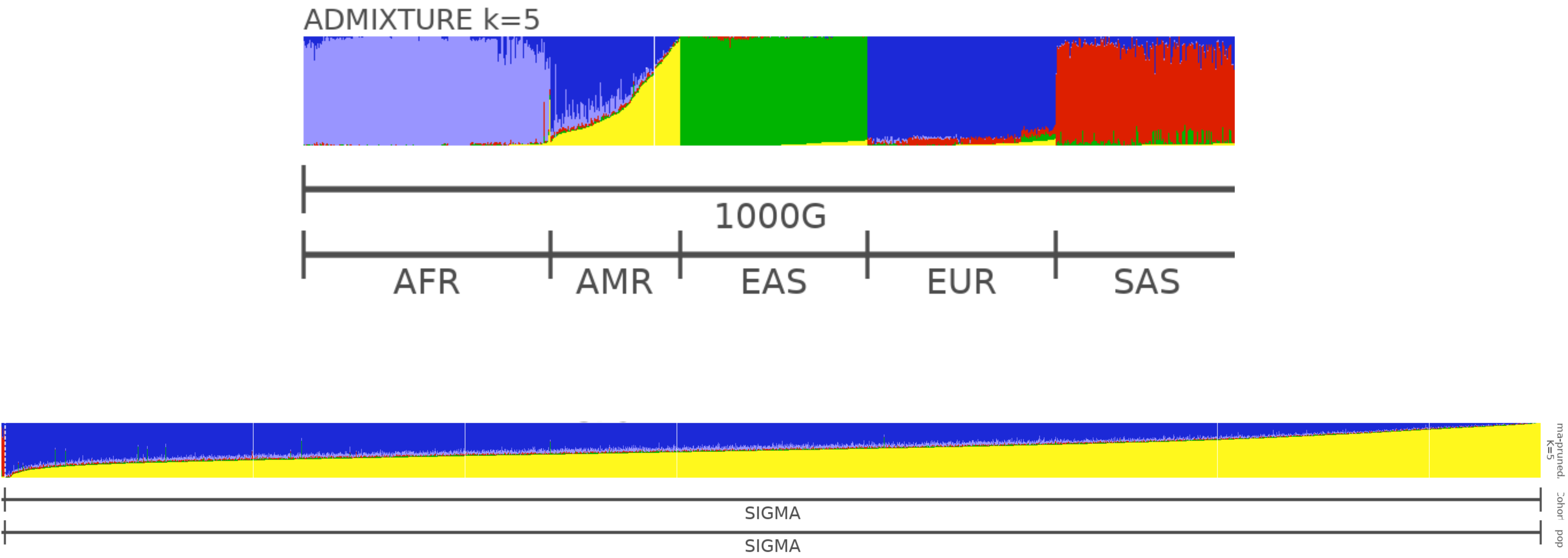


**S6 Fig. ADMIXTURE analysis (K=5) of individuals included in the SIGMA cohort and the 1000 Genomes Project.** Each individual is represented as a thin vertical bar. The colors can be interpreted as different ancestries. AFR represents African; AMR represents Admixed American; EAS represents East Asian; EUR represents European and SAS represents South Asian.

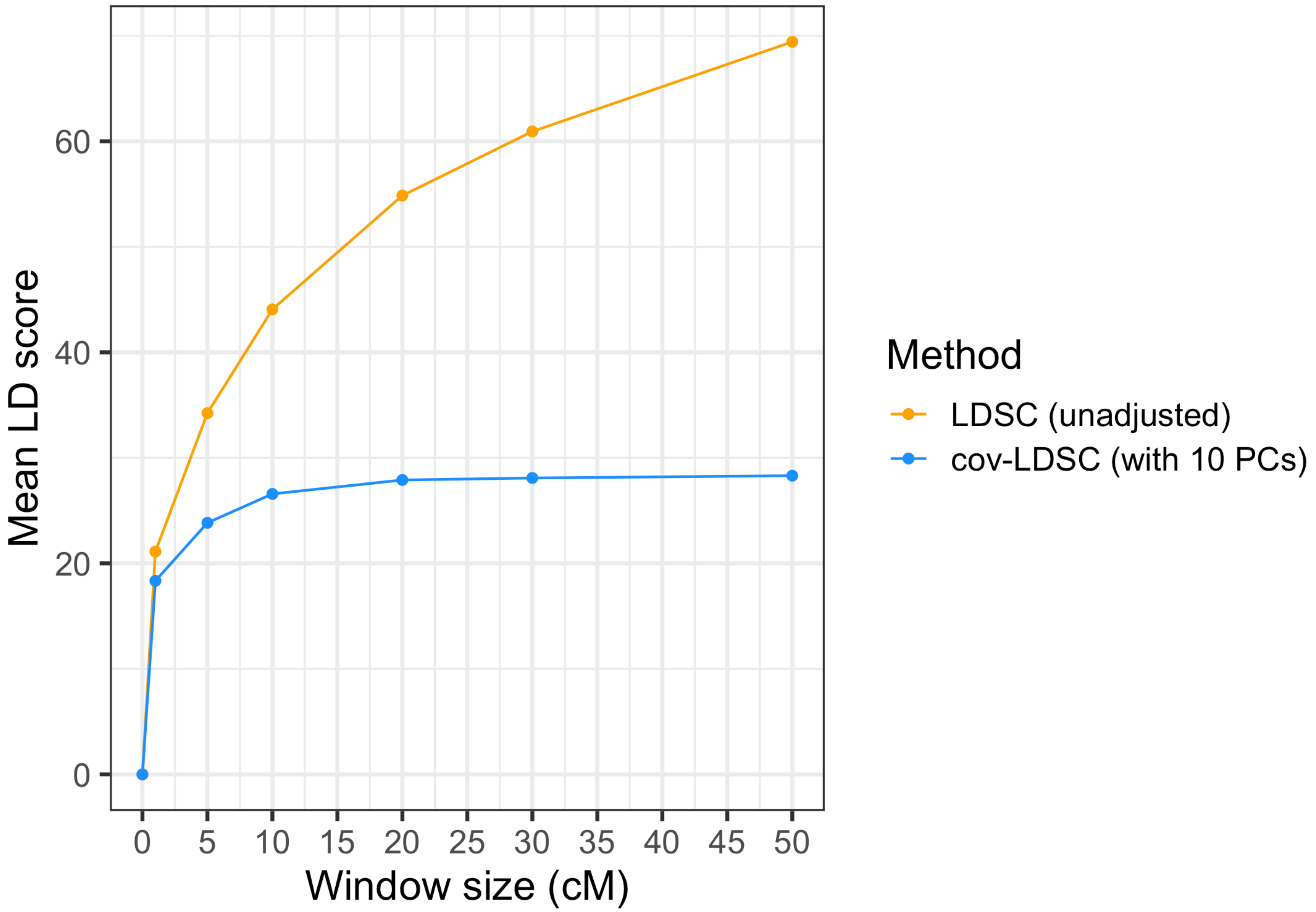


**S7 Fig. LD score estimates with varying window size in the SIGMA cohort.** LD score estimates using LDSC (orange) and cov-LDSC (blue) with varying window size in the SIGMA cohort (N=8,214). The x-axis shows the genomic window size used for estimating LD scores measured in centimorgan (cM). The y-axis shows the mean LD score estimates. For cov-LDSC, 10 PCs are used in all scenarios.

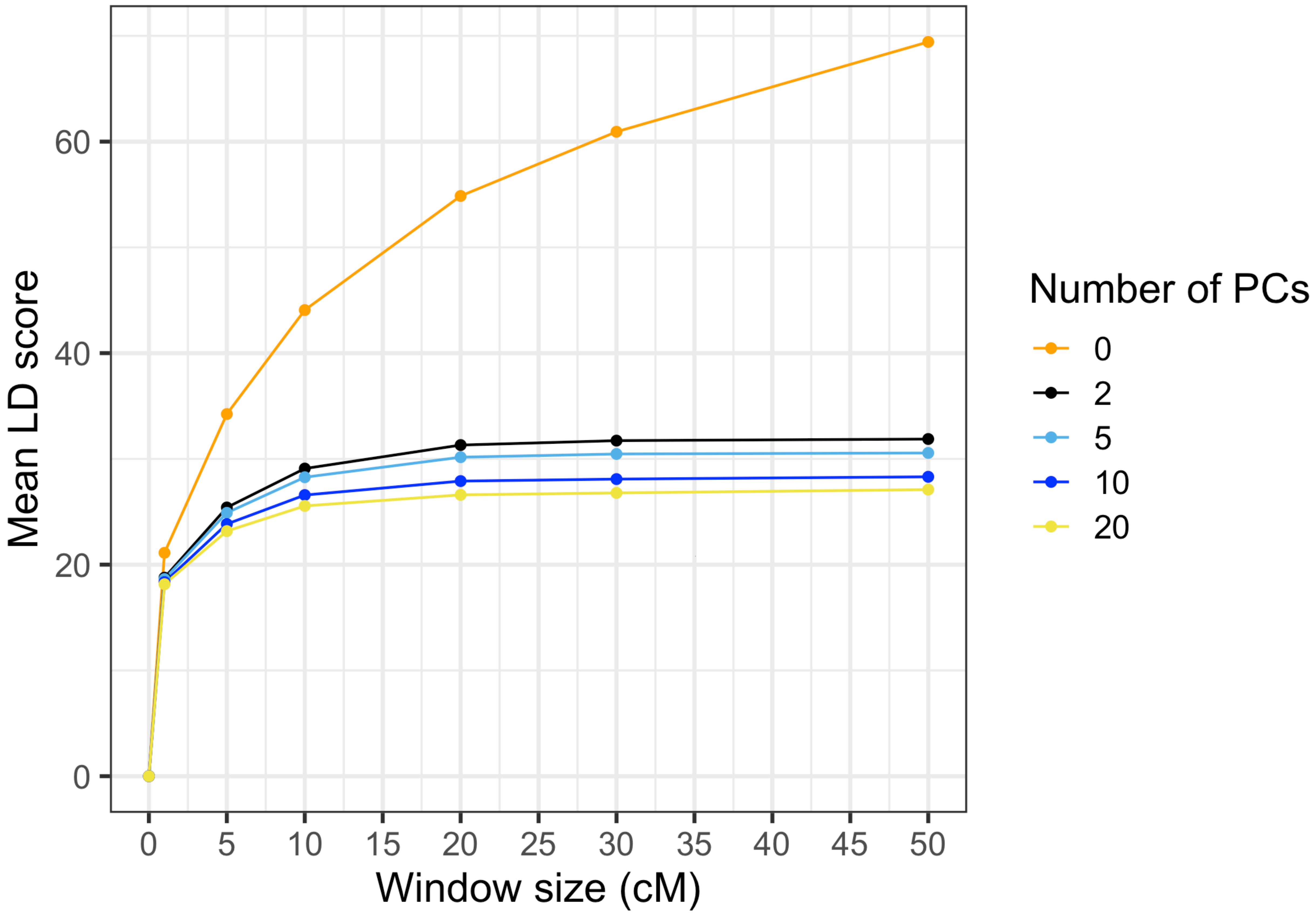


**S8 Fig. LD score estimates with varying window size and number of PCs in the SIGMA cohort.** LD score estimates (y-axis) using different number of PCs at different window sizes (x-axis).

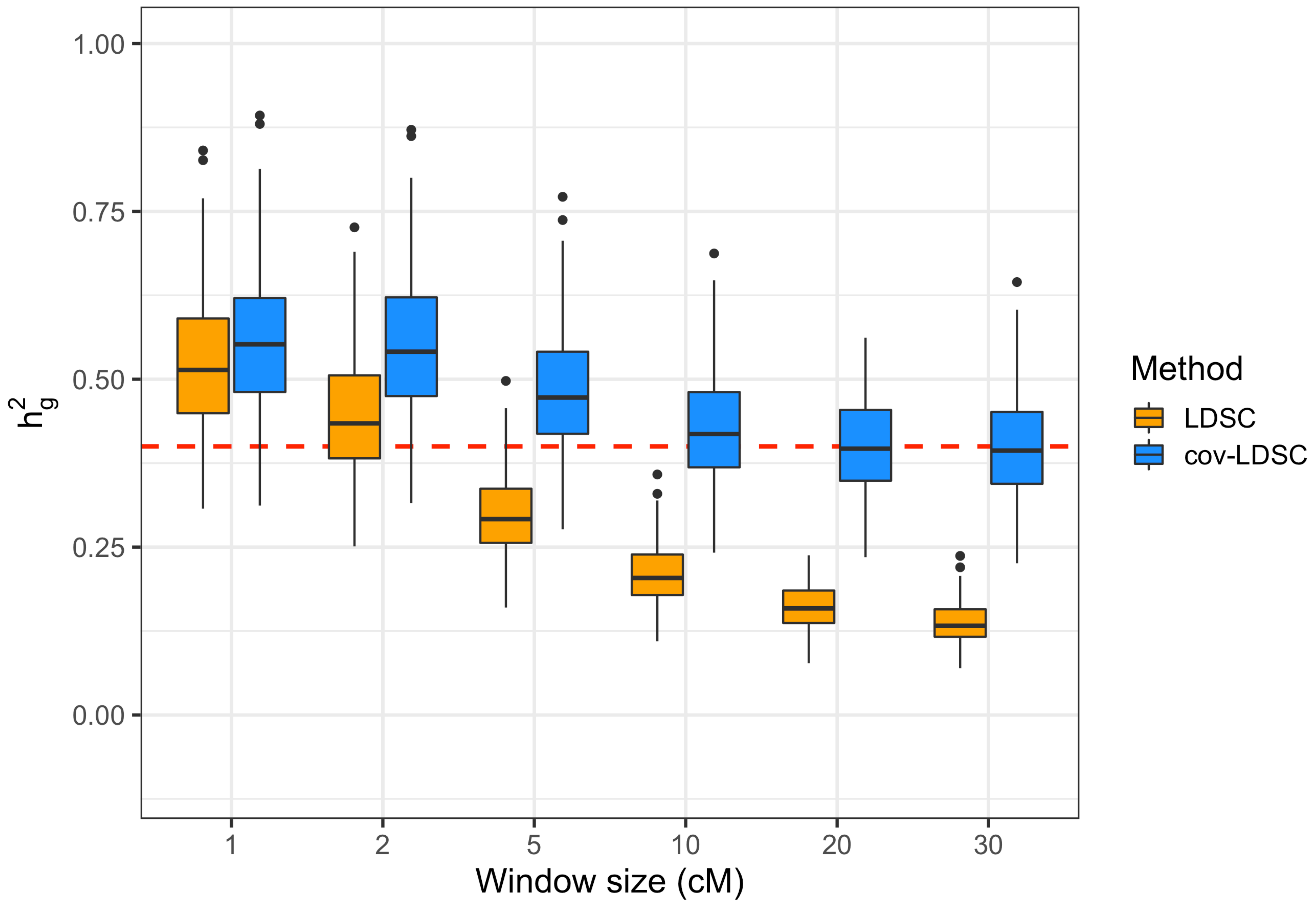


**S9 Fig. Estimates of heritability ( $h_g^2$ ) with varying window sizes used in LD score estimation in the SIGMA cohort.** cov-LDSC (blue) with 10 PCs and varying window size used to obtain LD score. We assumed a true $h_g^2$ of 0.4 and 1% causal variant in each simulation. 100 replicates are used for each window size.

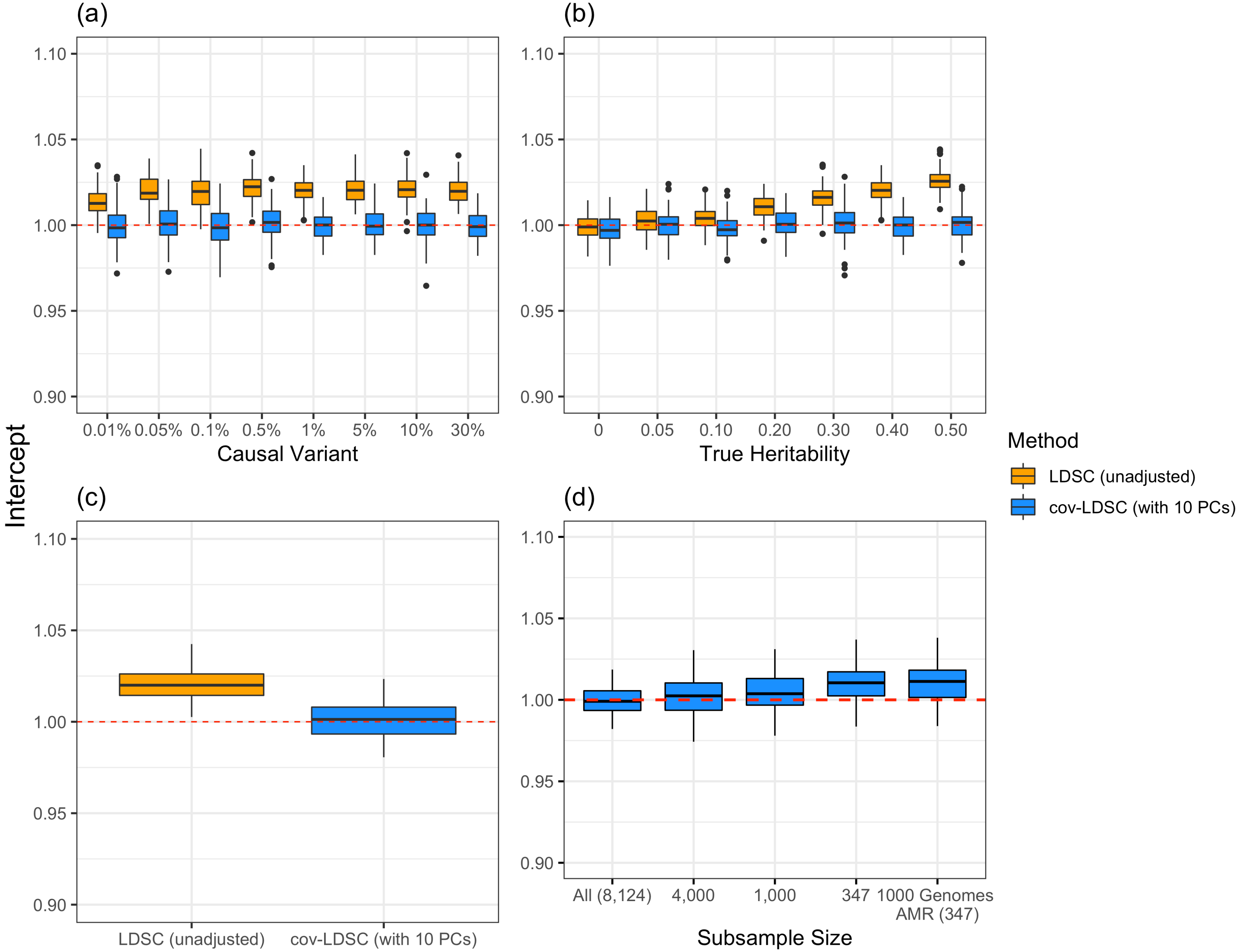


**S10 Fig. Intercept of estimated $h_g^2$ under different simulation scenarios using the SIGMA cohort as described in Figure 2.** LDSC (orange) underestimated $h_g^2$ and cov-LDSC (blue) yielded less biased $h_g^2$ estimates under all settings. Each boxplot represents the mean LD score estimate from 100 simulations of 8,124 individuals included in the SIGMA project. For cov-LDSC, a window size of 20-cM with 10 PCs are used in all scenarios.For LDSC, a window size of 20-cM are used in all scenarios A true polygenic quantitative trait with $h_g^2$=0.4 is assumed for scenarios (a), (c) and (d). 1% causal variants are assumed for scenarios (b)-(d). (a) Intercept with varying numbers of causal variants (0.01% - 50%). (b) Intercept with varying heritability (0, 0.05, 0.1, 0.2, 0.3, 0.4 and 0.5). (c) Intercept with the presence of an environmental stratification component aligned with the first PC of the genotype data is included in the phenotype simulation. (d) Intercept when using a subset of total samples and using admixed American samples included in the 1000 Genomes Project.

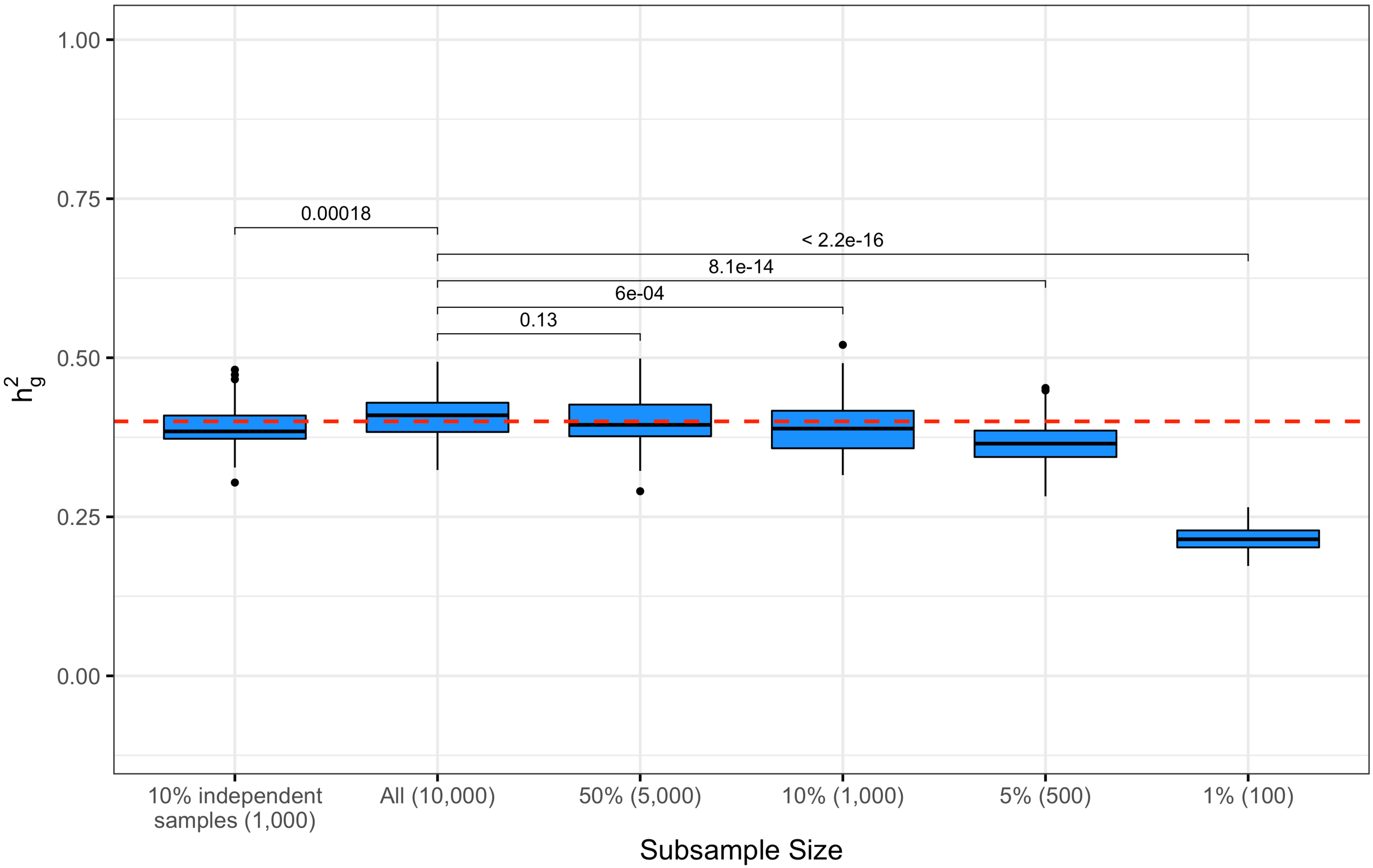


**S11 Fig. Estimates of heritability ($h_g^2$) in simulated genotypes using LD scores estimated with varying sample sizes.** cov-LDSC (blue) is used with varying sample sizes used to obtain LD scores. A random subset of 1%, 5%, 10% and 50% of the total samples (N=10,000) in the simulated genotypes are used to calculate *in-sample* LD scores and then to obtain hg2 estimates. LD scores are also obtained using independent genotypes (N=1,000) using the perfect matching demographic model.

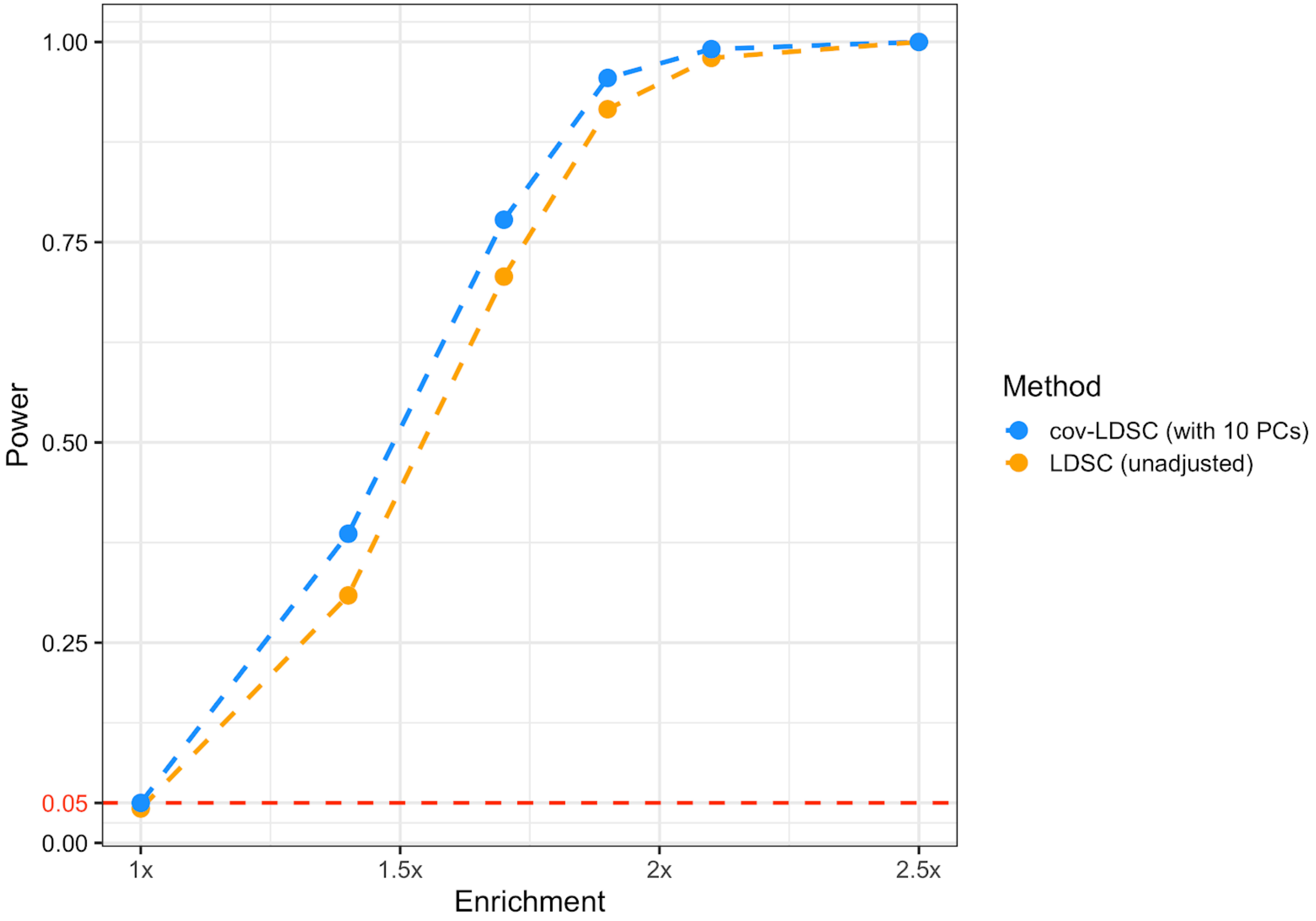


**S12 Fig. Simulation results assessing type I error and power for LDSC and cov-LDSC.** We simulated a polygenic trait with $h_g^2$ = 0.5. LDSC (orange) shows less power compared to cov-LDSC (blue) in detecting tissue. Each point shows the proportion of simulations (1,000 for each point) in which a null hypothesis of no tissue enrichment is rejected (Pr(rejected at P < 0.05)), as a function of the z-score of total SNP heritability.

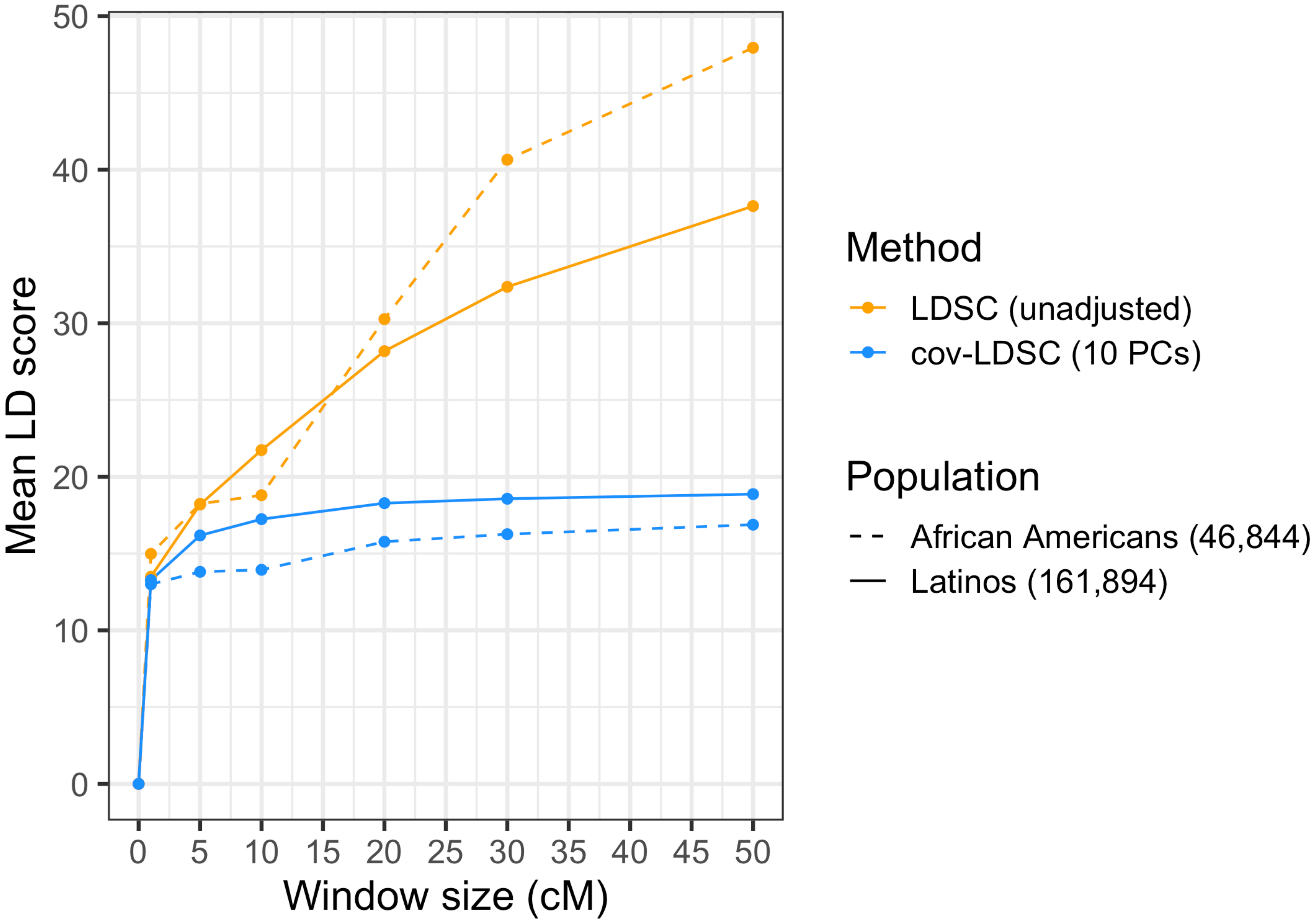


**S13 Fig. LD score estimates with varying window size in populations from 23andMe.** LD score estimates using unadjusted LDSC (orange) and cov-LDSC (blue) with 10 PCs with varying window size in both African Americans (N=46,844, dashed line) and Latinos (N=161,894, solid line) from the 23andMe cohort. The x-axis shows the genomic window size used for estimating LD scores measured in centimorgan (cM). The y-axis shows the mean LD score estimates.

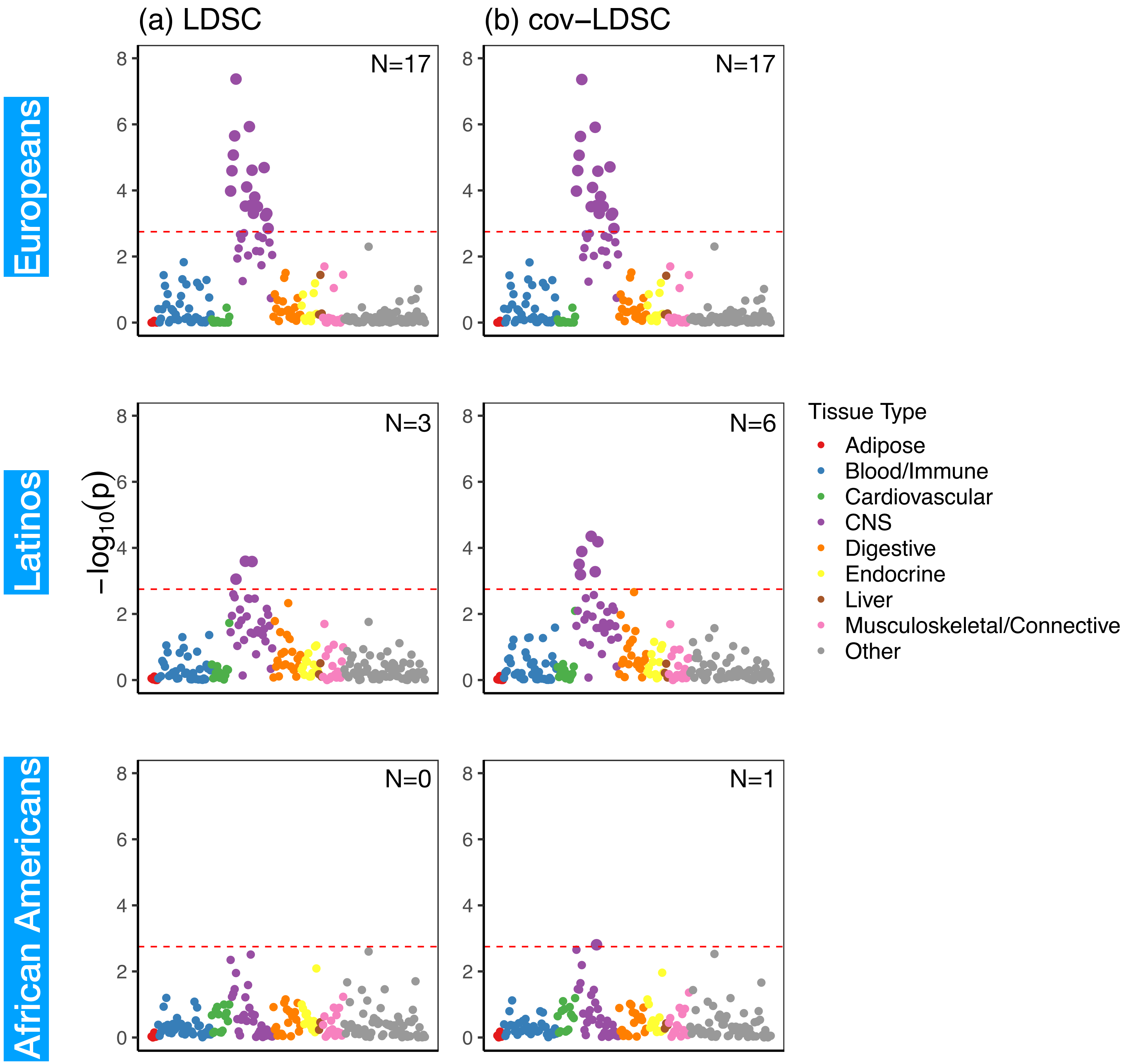


**S14 Fig. Tissue and cell type specific analysis with summary statistics in 23andMe Latinos using in-sample original LD and in-sample cov-LD for BMI.** The left panel (a) shows the tissue and cell type specific analysis using original LDSC with in-sample LD scores; while the right panel (b) shows the tissue and cell type specific analysis using cov-LDSC with in-sample cov-LD scores for BMI in 23andMe cohort. The label on the top right in each plot indicates the number of significant tissue type enrichments for each analysis. We observed no difference between LDSC and cov-LDSC in European populations. In contrast, we observed more enrichments and the p-values are more significant using cov-LDSC in Latinos and African Americans.

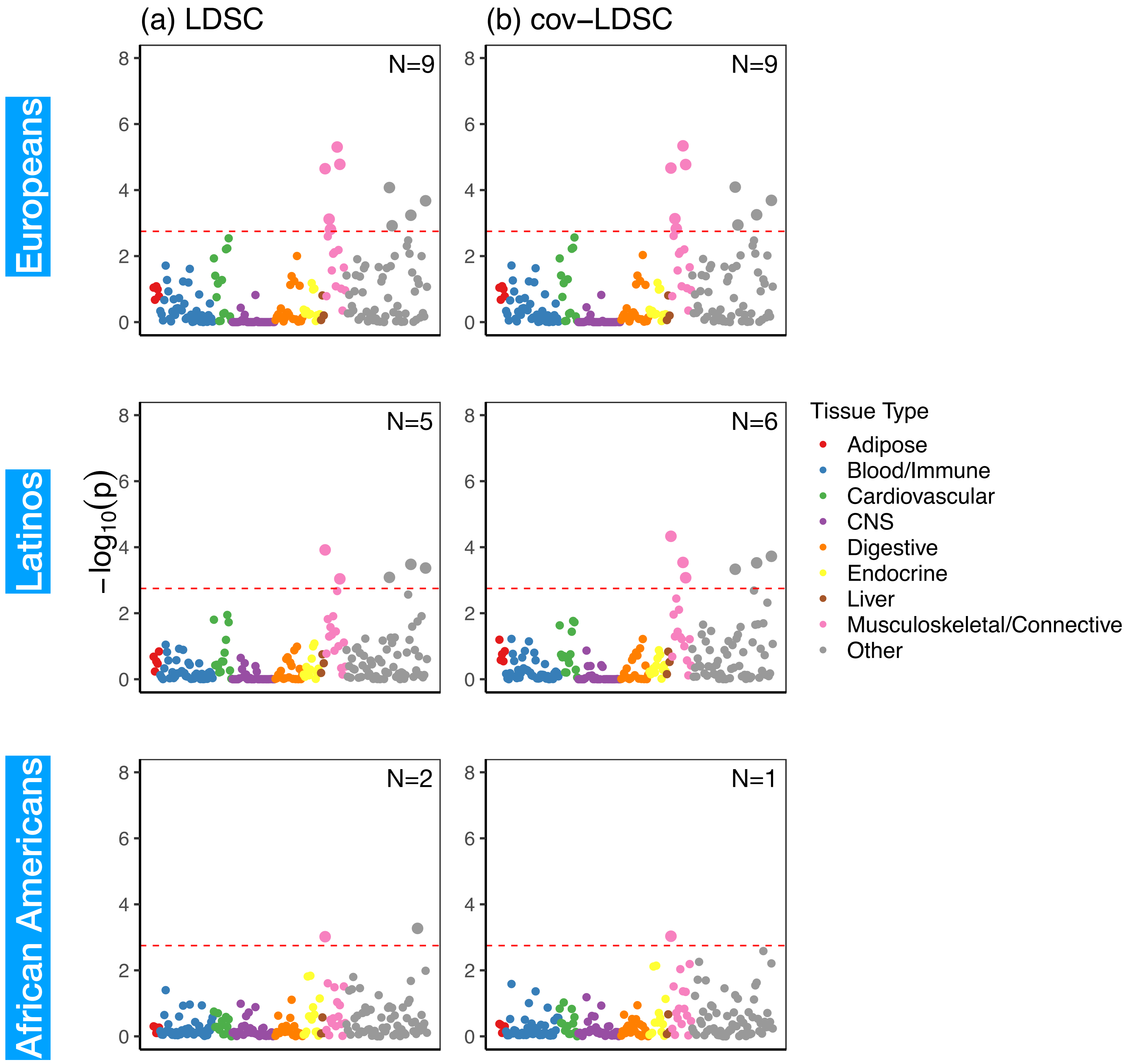


**S15 Fig. Tissue and cell type specific analysis with summary statistics in 23andMe Latinos using in-sample original LD and in-sample cov-LDSC for height.** The left panel (a) shows the tissue and cell type specific analysis using original LDSC with in-sample LD scores; while the right panel (b) shows the tissue and cell type specific analysis using cov-LDSC with in-sample cov-LD scores for BMI in 23andMe cohort. The label on the top right in each plot indicates the number of significant tissue type enrichments for each analysis. We observed no difference between LDSC and cov-LDSC in European populations. In contrast, we observed more enrichments and the p-values are more significant using cov-LDSC in Latinos and African Americans.

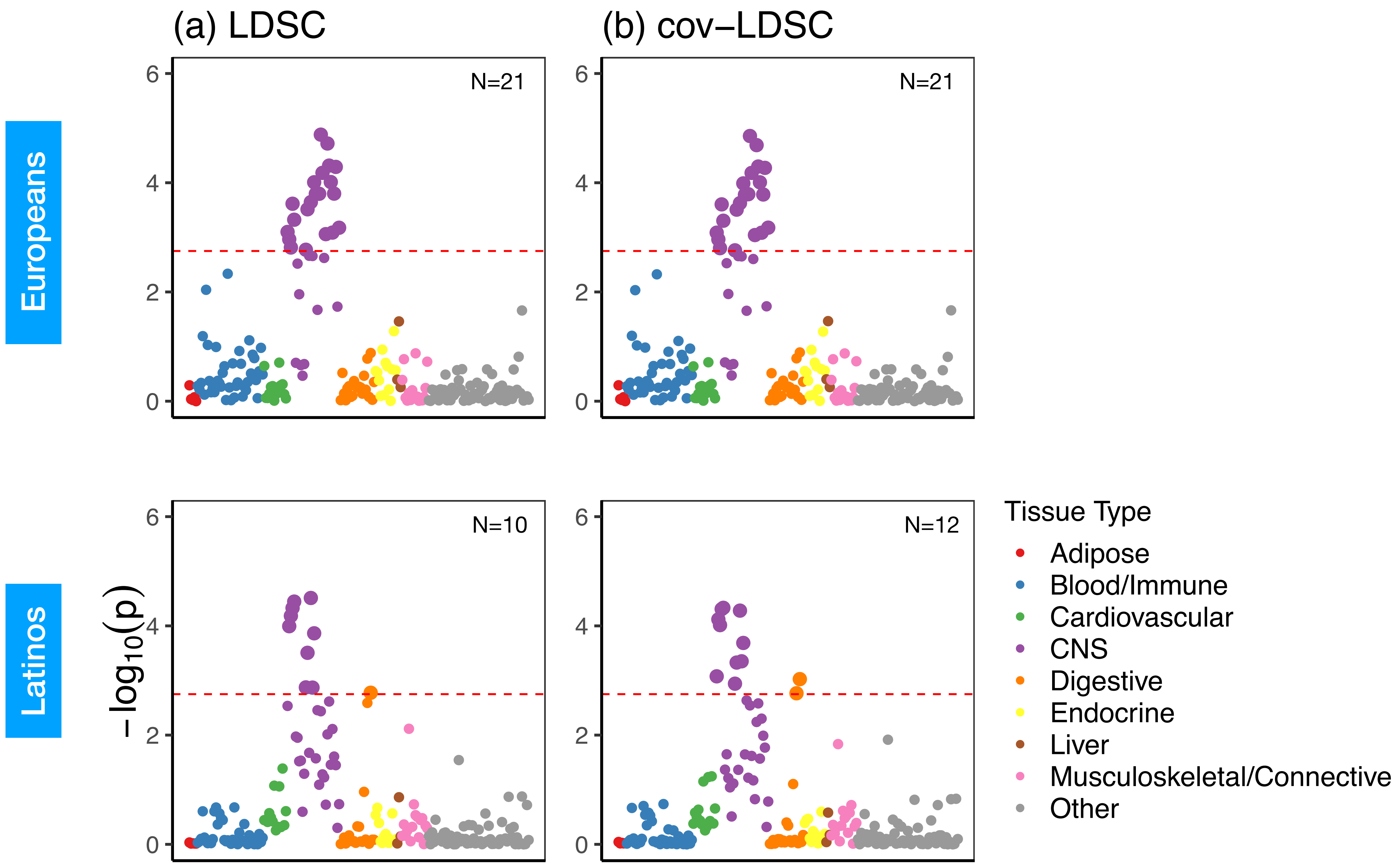


**S16 Fig. Tissue and cell type specific analysis with summary statistics in 23andMe Latinos using in-sample original LD and in-sample cov-LD for morning person.** The left panel (a) shows the tissue and cell type specific analysis using original LDSC with in-sample LD scores; while the right panel (b) shows the tissue and cell type specific analysis using cov-LDSC with in-sample cov-LD scores for BMI in 23andMe cohort. The label on the top right in each plot indicates the number of significant tissue type enrichments for each analysis. We observed no difference between LDSC and cov-LDSC in European populations. In contrast, we observed more enrichment in and around sets of genes that are specifically expressed in tissue- and cell-types using cov-LDSC in Latinos and African Americans.

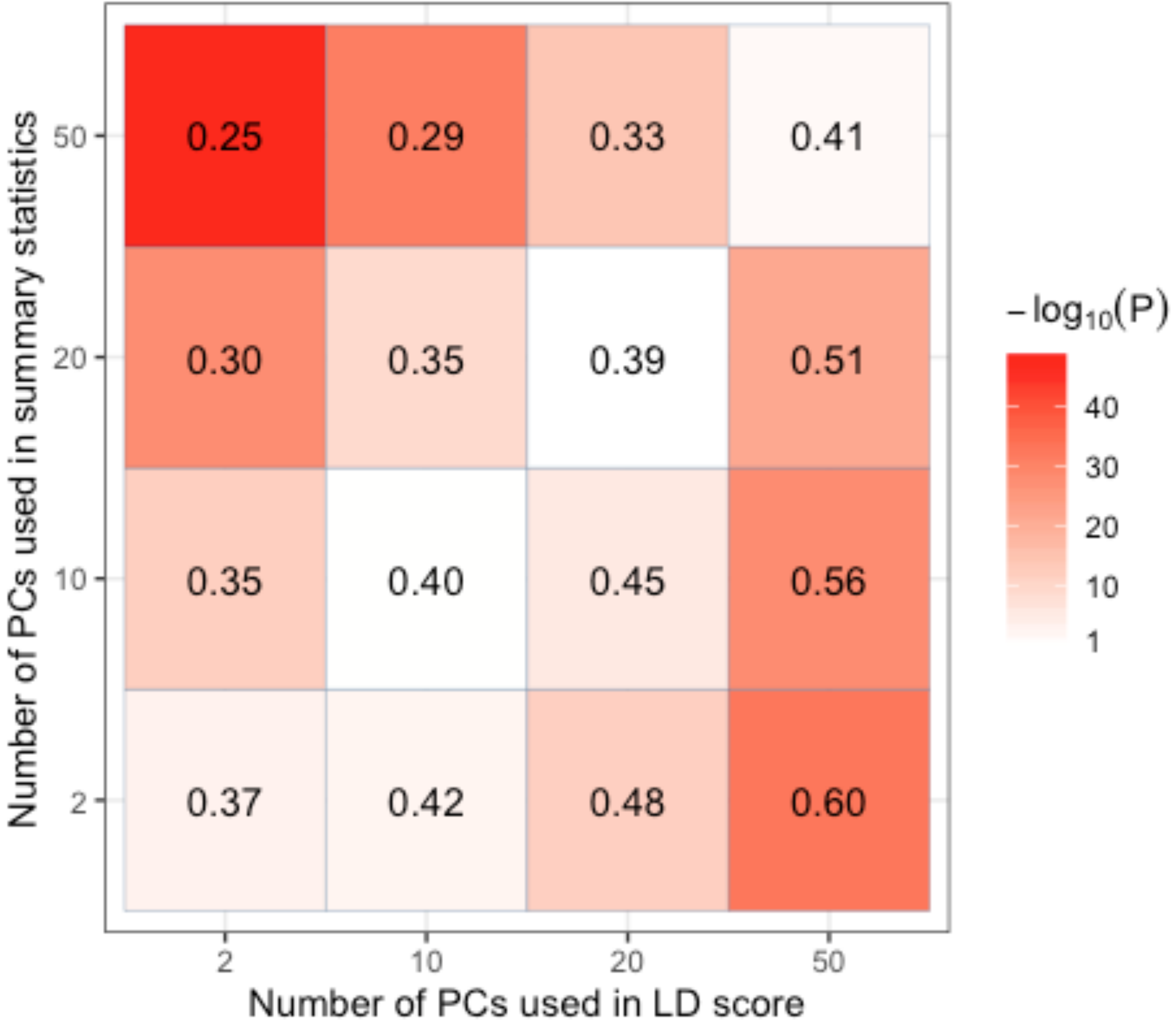


**S17 Fig. Heritability estimate with different number of PCs for GWAS association test and LD score adjustment.** We simulated the phenotypes on the SIGMA cohort using additive model assuming 1% causal SNPs with $h_g^2 = 0.4$. We performed univariate cov-LDSC to measure heritability. We varied number of PCs included in summary statistics and varied number of PCs used in cov-LDSC. The x-axis shows the number of PCs included in the cov-LDSC calculation and the y-axis shows the number of PCs included in the summary statistics calculation within the same sample. Numbers in each cell represent the mean $h_g^2$ estimates from 100 replications. The color (from white to red) represents the statistical difference between the estimated $h_g^2$ and the truth (measured in -log10(P)) . A red cell indicates the $h_g^2$ estimate is significantly different from the truth.

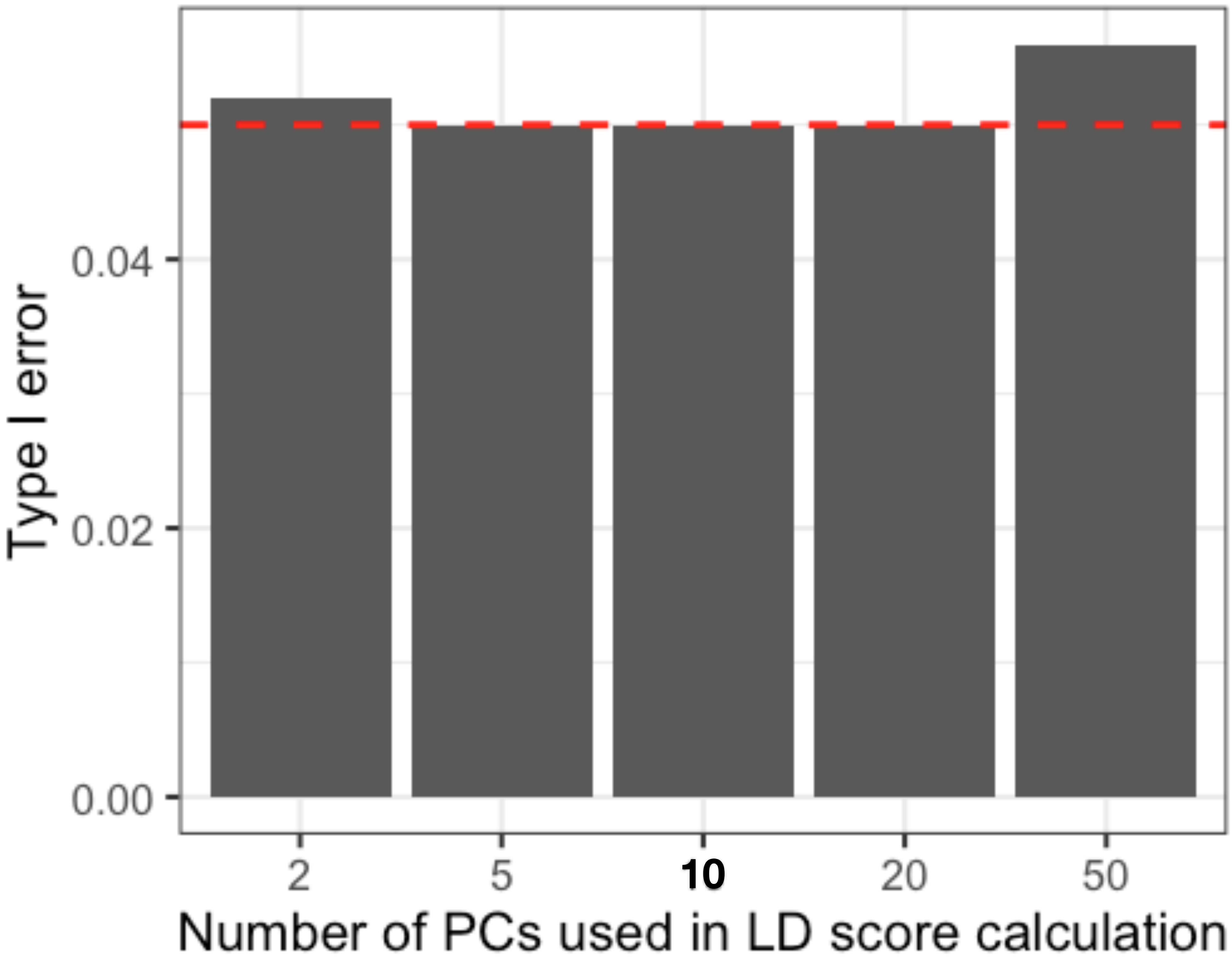


**S18 Fig. Type I error in tissue-type-specific enrichment when different number of PCs are used to generate summary statistics and LD scores.** We generated 1,000 simulations for scenarios where there are different number of PCs (2, 5, 10, 20 and 50) included when calculating LD scores and generating summary statistics (10 PCs) in the cell and tissue-specific enrichment analysis. We simulated a polygenic trait with $h_g^2 = 0.5$. Each bar shows the proportion of simulations in which a null hypothesis of no tissue enrichment is rejected (Pr(rejected at P < 0.05)), as a function of the z-score of total SNP heritability. The horizontal red line indicates P = 0.05.

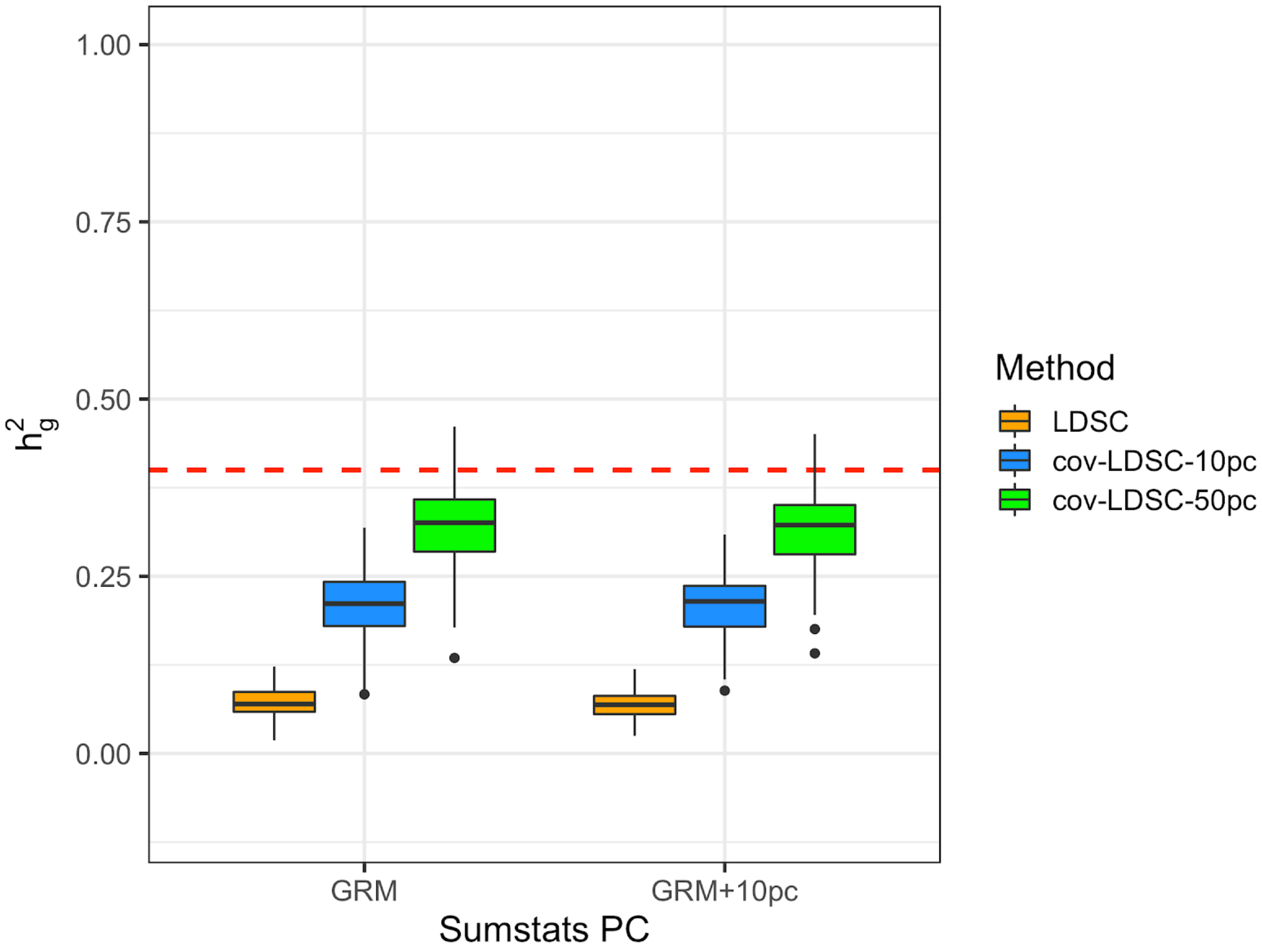


**S19 Fig. LDSC and cov-LDSC with summary statistics derived from linear mixed models.** Estimation of heritability (truth hg2= 0.4) using LDSC and cov-LDSC with 10 (blue) and 50 (green) PCs and a window size of 20-cM. Each boxplot represents the mean LD score estimate from 100 simulations of genotypes from the 8,124 individuals included in the SIGMA cohort. All summary statistics are derived from linear mixed models with genetic relationship matrix (GRM) only or GRM with 10 genome-wide PCs using GEMMA.

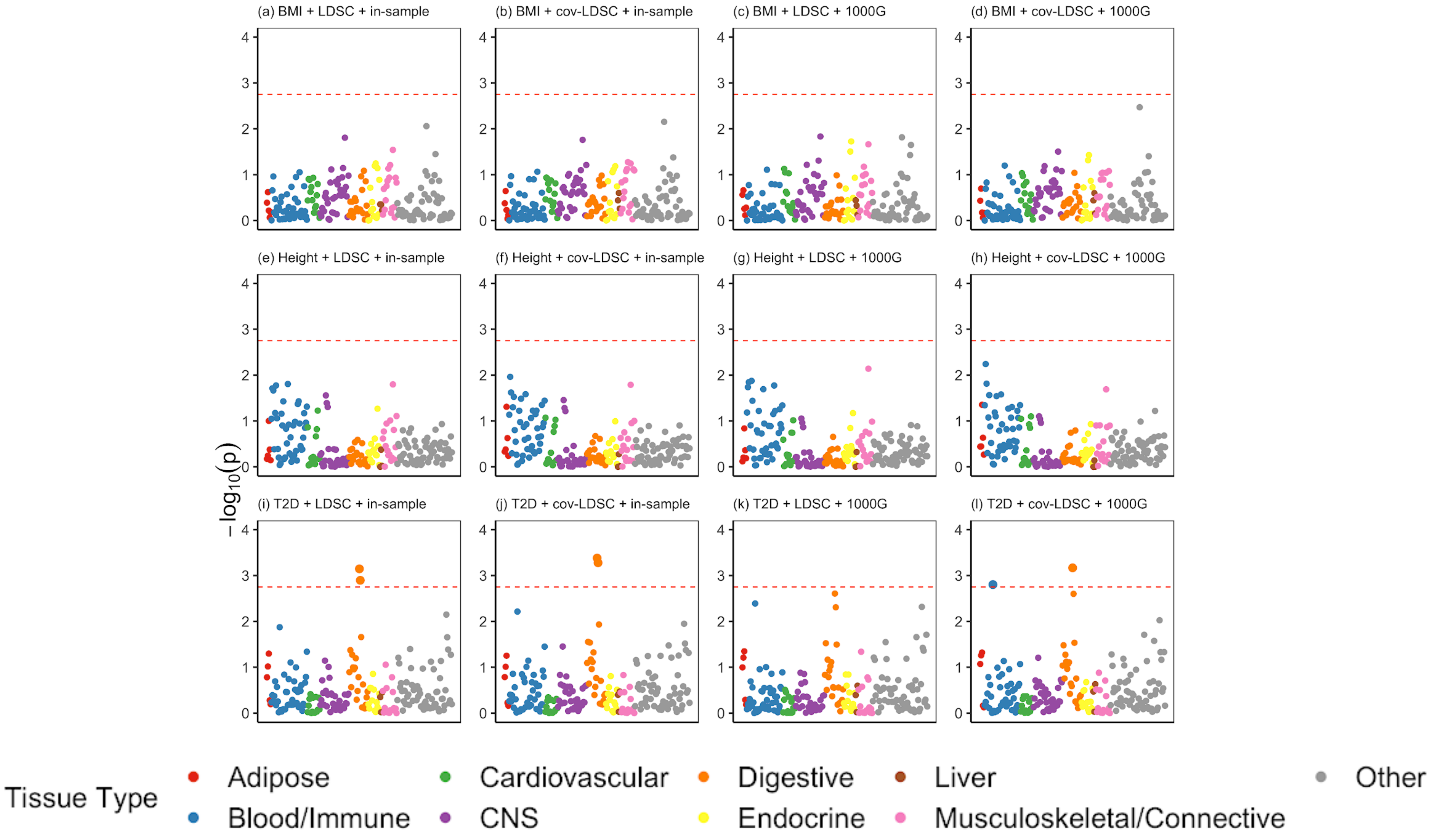


**S20 Fig. Results of multiple-tissue analysis for body mass index (BMI), height and type 2 diabetes (T2D) in the SIGMA cohort using in-sample and out-of-sample LD panels using LDSC and cov-LDSC, respectively.** Each point represents a tissue type from either the GTEx data set or the Franke lab data(33,34). From left to right, (a-d) show multiple-tissue analysis for BMI, when using LDSC and cov-LDSC with in-sample and out-of-sample LD reference panels. (e-h) show multiple-tissue analysis for height (e-h) when using LDSC and cov-LDSC with in-sample and out-of-sample LD reference panels. (i-l) show multiple-tissue analysis for T2D when using LDSC and cov-LDSC with in-sample and out-of-sample LD reference panels.

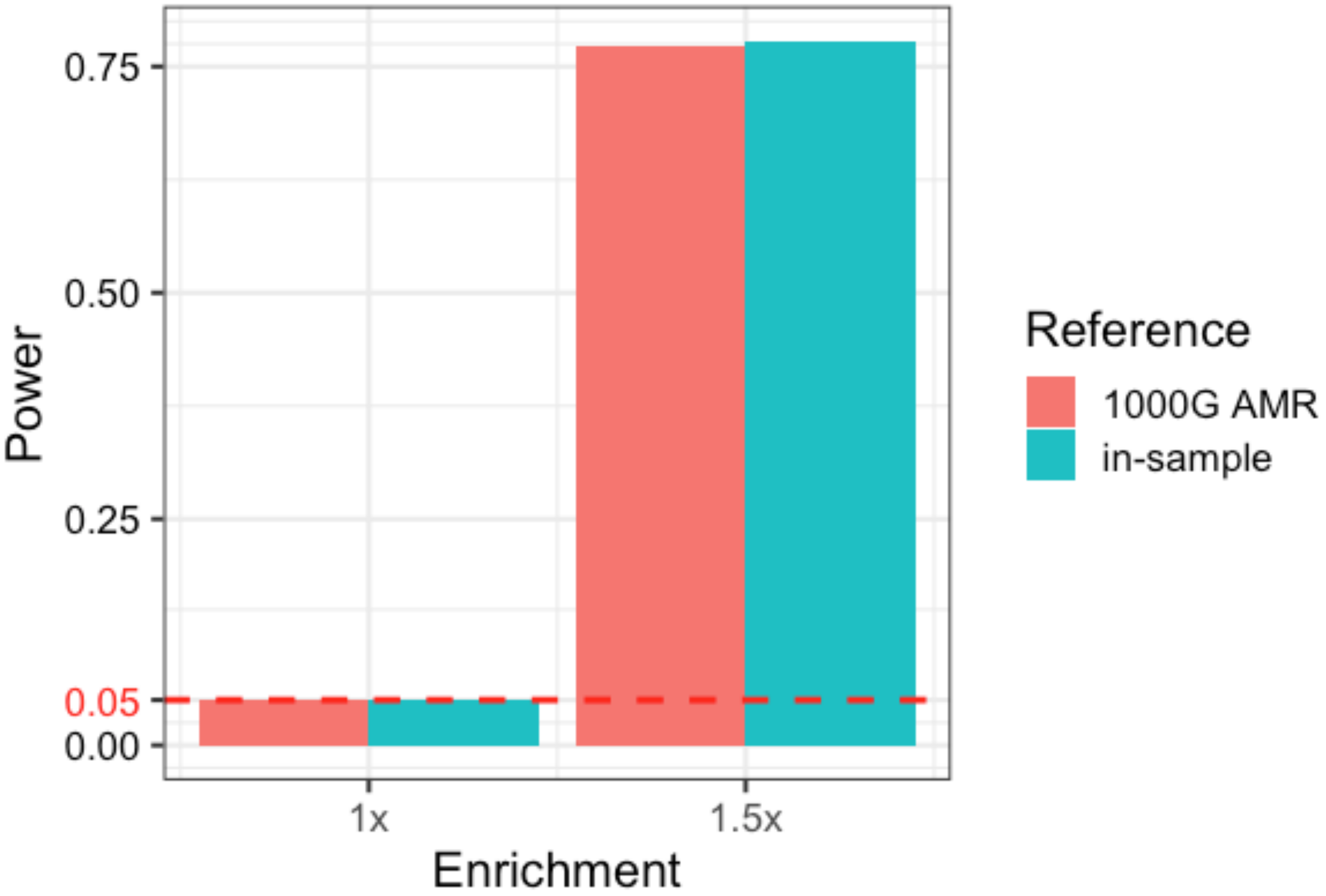


**S21 Fig. Enrichment analysis using in-sample and out-of-sample LD reference panel.** We simulated a polygenic trait with $h_g^2 = 0.5$. Similar power was obtained when using in-sample (obtained from the SIGMA cohort, turquoise) and out-of-sample (obtained from 1000 Genomes Admixed American (AMR) samples, red) reference panel. In both cases, type I error (at no (1x) enrichment) are well controlled. Each bar shows the proportion of simulations (1,000 for each point) in which a null hypothesis of no tissue enrichment is rejected (Pr(rejected at P < 0.05), red horizontal line), as a function of the z-score of total SNP heritability.

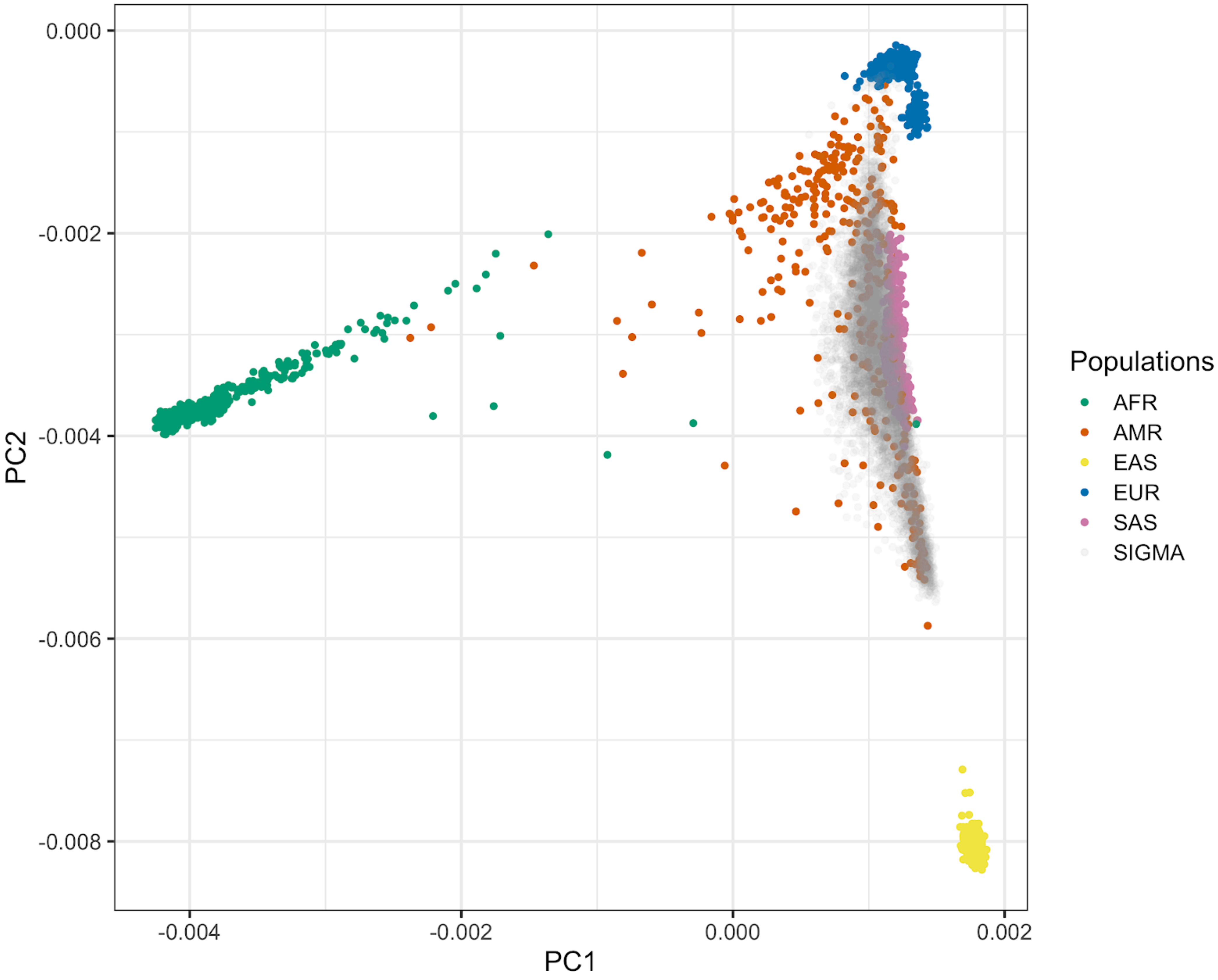


**S22 Fig. Principal component analysis (PCA) of the SIGMA samples.** Samples included in the SIGMA cohort projected onto the first two principal components using SNP weights precomputed from samples in the 1000 Genomes Phase 3 project using SNP weights. AFR represents Africans (green); AMR represents Admixed Americans (orange); EAS represents East Asians (yellow); EUR represents Europeans (blue); SAS represents South Asians (pink) and SIGMA samples are presented in gray.

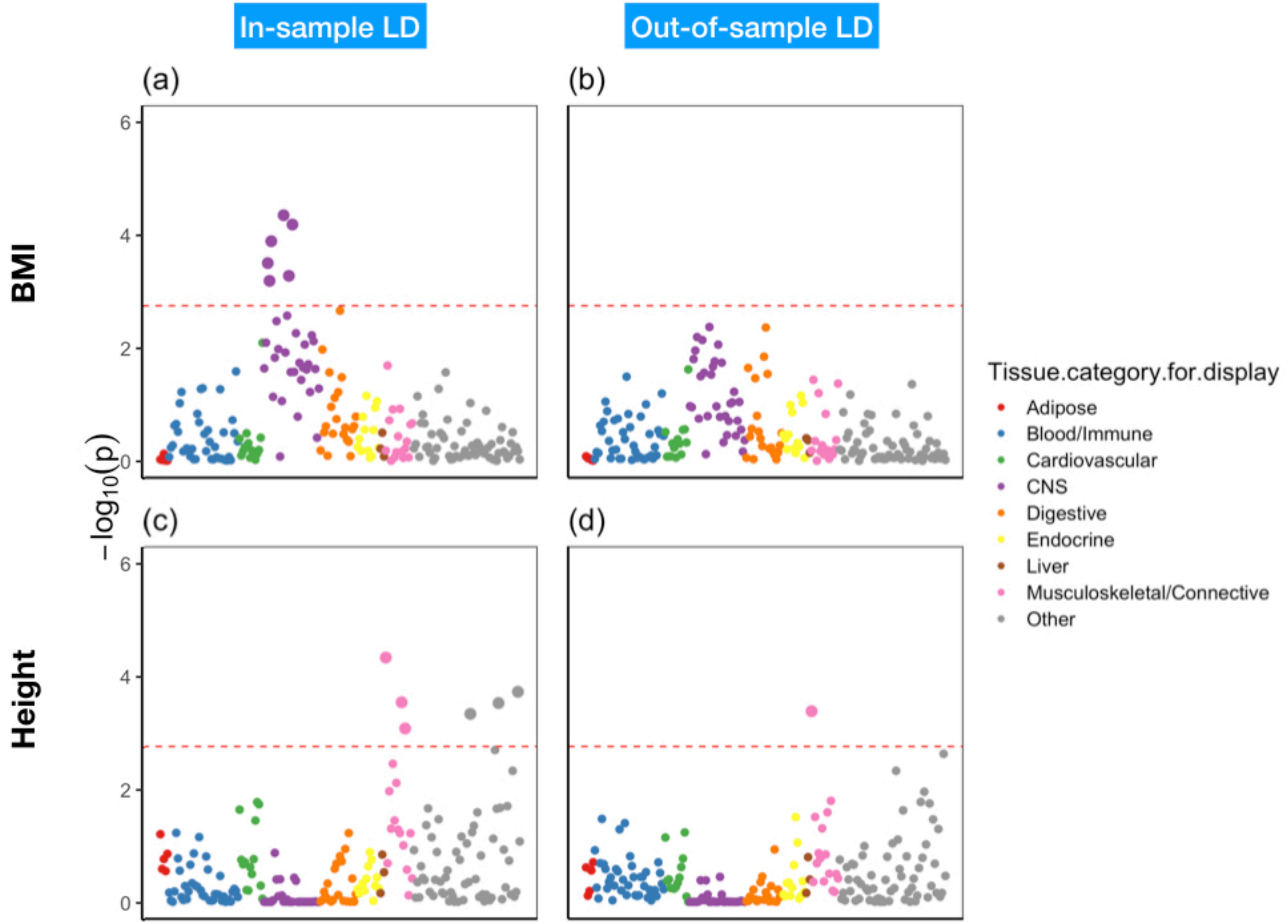


**S23 Fig. Tissue and cell type specific analysis with summary statistics in 23andMe Latinos using in-sample cov-LD and out-of-sample cov-LD obtained using 1000G AMR samples.** In sample LD is obtained in 23andMe Latinos with 20cM window size and 10PCs. We observed cell type enrichments in both BMI and height using in-sample cov-LD. However, when we used out of sample 1000G AMR cov-LD with 20cM window size and 10PCs, we observed no cell type enrichments in either BMI and height.

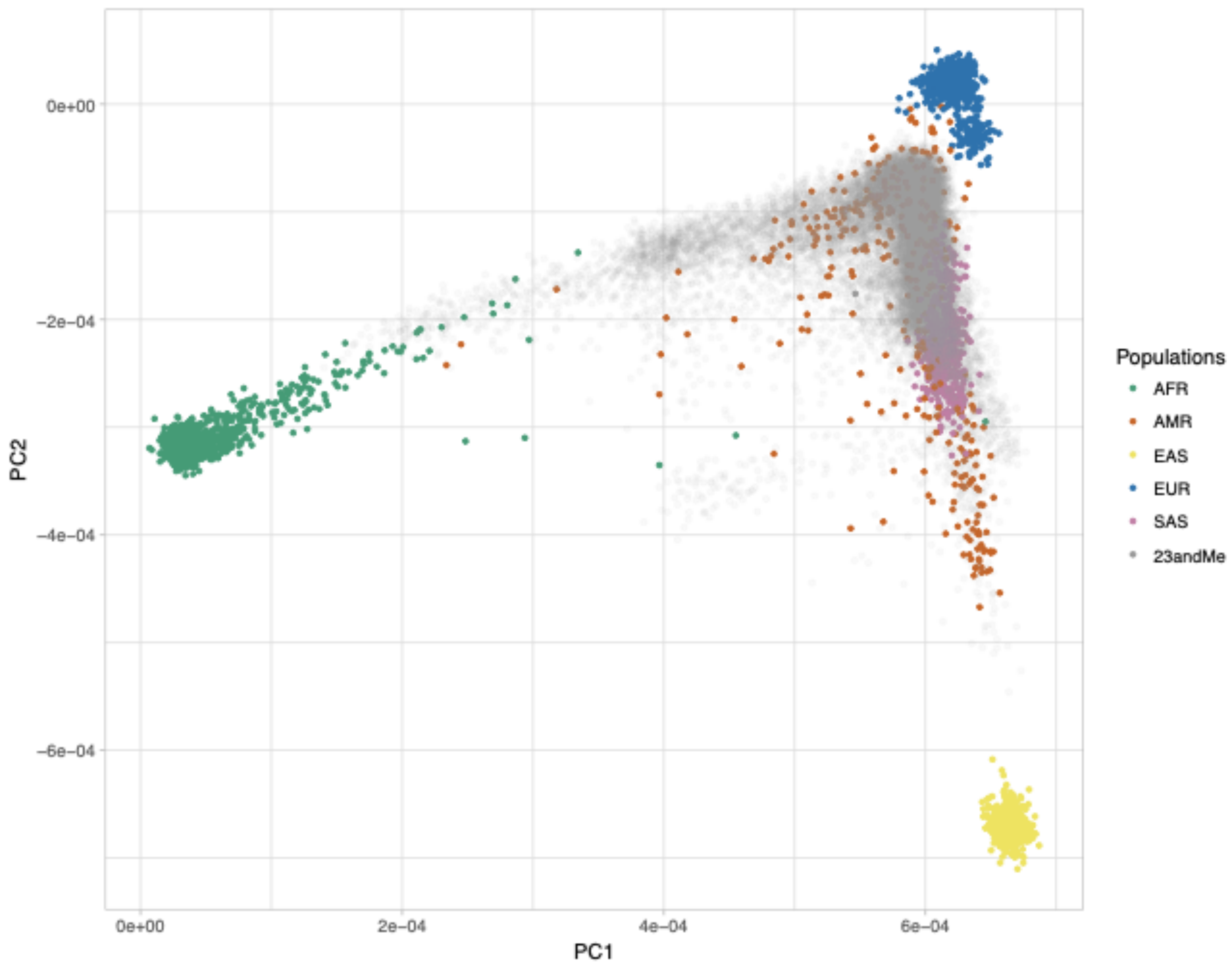


**S24 Fig. Principal component analysis (PCA) of the 23andMe samples.** Samples included in the SIGMA cohort projected onto the first two principal components using SNP weights precomputed from samples in the 1000 Genomes Phase 3 project using SNP weights. AFR represents Africans (green); AMR represents Admixed Americans (orange); EAS represents East Asians (yellow); EUR represents Europeans (blue); SAS represents South Asians (pink) and SIGMA samples are presented in gray.
